# The exogenous application of the apocarotenoid retinaldehyde negatively regulates auxin-mediated root growth

**DOI:** 10.1101/2023.09.12.555306

**Authors:** Kang Xu, Haoran Zeng, Feiyang Lin, Emi Yumoto, Masashi Asahina, Ken-ichiro Hayashi, Hidehiro Fukaki, Hisashi Ito, Masaaki K. Watahiki

**Affiliations:** Graduate School of Life Science, Hokkaido University, Sapporo, Japan; Advanced Instrumental Analysis Center. Teikyo University, Utsunomiya, Japan; Department of Biosciences, Teikyo University, Utsunomiya, Japan; Department of Bioscience, Okayama University of Science, Okayama, Japan; Department of Biology, Graduate School of Science, Kobe University, Kobe, Japan; Institute of Low Temperature Science, Hokkaido University, Sapporo, Japan; Division of Biological Sciences, Faculty of Science, Hokkaido University, Sapporo, Japan

**Author notes:** **List of authors:** Kang Xu; Haoran Zeng; Feiyang Lin; Emi Yumoto; Masashi Asahina; Ken-ichiro Hayashi; Hidehiro Fukaki; Hisashi Ito; Masaaki K. Watahiki. The author responsible for distribution of materials integral to the findings represented in this article in accordance with the policy described in the Instruction for Authors (www.plantphysiol.org) is Masaaki K. Watahiki. ResearcherID: GFY-7778-2022.

## Abstract

Root development is essential for plant survival. The lack of carotenoid biosynthesis in the *phytoene desaturase 3* (*pds3*) mutant results in short primary roots (PR) and reduced lateral root (LR) formation. In this study, we show that short-term inhibition of PDS by fluridone suppressed PR growth in WT, but to a lesser extent in auxin mutants. Such an inhibition of PDS activity increased endogenous indole-3-acetic acid (IAA) levels, promoted auxin signaling, and partially complemented the PR growth of auxin deficient mutant, *YUCCA 3 5 7 8 9* quadruple mutant (*yucQ*). The exogenous application of retinaldehyde (retinal), an apocarotenoid derived from β-carotene, complemented the fluridone-induced suppression of root growth, as well as the short roots of the *pds3* mutant. Retinal also partially complemented the auxin-induced suppression of root growth. These results suggest that retinal may play a role in regulating root growth by modulating endogenous auxin levels.

**One-sentence summary:** The short-term inhibition of carotenoid biosynthesis mediates the production of β-carotene-derived apocarotenoid retinal which negatively regulates auxin levels and signaling to control root growth.

## Introduction

The root system plays a crucial role in water and nutrient uptake, anchoring plants to the soil, and serving as a source of signaling chemicals. Primary root (PR) elongation and lateral root (LR) branching are controlled by environmental cues and determine the architecture of the root system, allowing plants to adapt to the environment. The phytohormone auxin controls root growth and LR patterning (Moreno-Risueno et al., 2010; Saini et al., 2013; Chen et al., 2014). An early auxin signaling requires a family of repressor genes known as *AUXIN/INDOLE-3-ACETIC ACID* (*Aux/IAA*s). *Aux/IAA* dimerizes with the transcription factor *AUXIN RESPONSE FACTOR* (*ARF*) and suppresses the transcriptional function of *ARF*s (Leyser, 2002; De Smet et al., 2010). Application of exogenous auxin suppresses PR length and induces LR formation (Hobbie and Estelle 1994; Okumura et al., 2013). Overexpression of *ARF19* reduces PR length, suggesting a negative role for auxin signaling in PR elongation (Okushima et al., 2005). Polar auxin transport (PAT), which controls the flow of auxin to create an auxin gradient, is also involved in root growth and LR formation (Reed et al., 1998; Vieten et al., 2007). The auxin permease mutant, *auxin resistant 1* (*aux1*) exhibits both agravitropism and resistance to PR growth suppression by exogenous IAA (Yamamoto and Yamamoto, 1998). The longer PRs of *aux1* compared to wild-type (WT) also suggest a negative role of endogenous auxin in PR growth (Roman et al., 1995).

*YUCCA* (*YUC*) encodes flavin monooxygenases which converts indole-3-pyruvic acid (IPyA) to indole-3-acetic acid (IAA) (Won et al., 2011; Zhao, 2012). *YUCCA* consists of a family of 11 genes and is highly redundant among them (Cheng et al., 2006; Xu et al., 2017). *YUC3*, *5*, *7*, *8*, and *9* are predominantly expressed in roots, and their quintuple mutant *yucQ* has a shorter PR and about half the amount of free IAA in roots compared to WT (Chen et al., 2014). The root meristem of *yucQ* is smaller and partially differentiated (Chen et al., 2014), suggesting that an appropriate level of auxin is required to maintain root meristem activity. Anthranilate is a precursor of tryptophan biosynthesis and contributes to IAA biosynthesis. The *WEAK ETHYLENE INSENSITIVE 2/ANTHRANILATE SYNTHASE α1* (*WEI2/ASA1*) and *WEI7/ANTHRANILATE SYNTHASE β1* (*WEI7/ASB1*) genes encode the subunits of anthranilate synthase, and their double mutant, *wei2 wei7* exhibits a dwarf phenotype with severely reduced PR length and IAA levels (Stepanova et al., 2005; Doyle et al., 2019).

Auxin regulates not only root development but also chloroplast development and chlorophyll accumulation. The *auxin resistant 1* (*axr1*) and *massugu1* (*msg1*)/*arf7* mutant, which are insensitive to auxin, are resistant to leaf chlorosis caused by 2,4-dichlorophenoxyacetic acid (2,4-D) (Watahiki and Yamamoto, 1997). The auxin signaling genes, *SOLITARY-ROOT/AUXIN/INDOLE-3-ACETIC ACID14* (*SLR/AUX/IAA14*), *ARF7*, and *ARF19*, are coupled with light and cytokinin signaling to regulate the gene sets associated with chloroplast development in the root (Kobayashi et al., 2012). Carotenoids are a diverse group of plastidial isoprenoid pigments that are essential for photosynthesis (Ruiz-Sola and Rodríguez-Concepción, 2012). In flowering plants, carotenoids are synthesized in the 2-C-methyl-D-erythritol 4-phosphate/1-deoxy-D-xylulose 5-phosphate (MEP/DOXP) pathway (Ruiz-Sola and Rodríguez-Concepción, 2012). PHYTOENE SYNTHASE (PSY) and PHYTOENE DESATURASE (PDS) are enzymes that convert geranylgeranyl pyrophosphate (GGPP) into phytoene and phytoene into ζ-carotene, respectively (Breitenbach and Sandmann, 2005; Rodríguez-Villalón et al., 2009). ζ-carotene then undergoes desaturation, isomerization and cyclization to generate carotenes like α-carotene and β-carotene (Ruiz-Sola and Rodríguez-Concepción, 2012). Carotenoids act not only as photopigments but also as precursors of phytohormones, abscisic acid (ABA) and strigolactones (SL) (Nambara and Marion-Poll, 2005; Umehara et al., 2008). Besides, the discovery of new bioactive apocarotenoids, which are the products of the oxidative cleavage of carotenoids, has increased during this century (Auldridge et al., 2006; Zheng et al., 2021). β-cyclocitral is formed by the oxidative cleavage of the C7’, C8’ double bond position of β-carotene and promotes PR growth and LR formation (Rodrigo et al., 2013; Dickinson et al., 2019). Anchorene, formed by cutting the C11, C12 and C11’, C12’ bonds in almost all plant carotenoids, promotes the adventitious root formation by modulating auxin distribution (Jia et al., 2019). Impairment of apocarotenoid biosynthesis shows shorter roots and reduced LR formation compared to WT (Van Norman et al., 2014). This suggests a role for carotenoids and apocarotenoids in root growth regulation. The apocarotenoid retinaldehyde (retinal), a precursor of vitamin A and retinoic acid in vertebrates formed by the oxidative cleavage of the C15, C15’ bond of β-carotene (Olson and Hayaishi, 1965), is mysteriously formed in flowering plants (Koschmieder et al., 2021), and is found to play a role in the periodic formation of LR (Dickinson et al., 2021).

In this study, we found that the herbicide fluridone (Bartels and Watson, 1978), which inhibits the enzymatic activity of PDS, suppressed root growth and partially mimicked the dwarf phenotype of the *pds3* mutant (Qin et al., 2007). This mechanism of root growth suppression remains to be elucidated. Here, we show that fluridone rapidly induces endogenous auxin levels and activates the auxin response. Genetic analysis revealed that this rapid response is dependent on PDS activity. Complementary approaches revealed that the apocarotenoid retinal negatively regulates endogenous auxin levels. Our results suggest that PDS activity positively regulates auxin catabolism to maintain auxin homeostasis via retinal.

## Results

### Fluridone suppresses the growth of the primary root

After transferring 5-day-old seedlings to fluridone medium or mock medium, the position of the root tip was marked and incubated for another 4 days. Fluridone suppressed PR growth in a dose-dependent manner (**Fig. 1, B and C**), and bleached the newly emerged true leaves, while the established cotyledons exhibited chlorosis (Supplemental Fig. S4A). The LR number and density in the newly grown areas of PRs after transfer to fluridone medium and the pre-grown areas of PRs were measured. The LR number in the pre-grown areas was induced by fluridone. In contrast, fluridone reduced the number of LRs in the newly grown areas (Supplemental Fig. S3). Overall, LR density was increased by fluridone (**Fig. 1D**). Other PDS inhibitors, such as norflurazon, diflufenican and picolinafen, also suppressed PR growth and increased LR density (Supplemental Fig. S2). To investigate the target specificity of fluridone, a fluridone-insensitive trait of PDS (*35S::mHvPDS*) (Arias et al., 2006) was then introduced into the *Arabidopsis* plants. The *35S::mHvPDS* plants were confirmed to have resistance to fluridone in terms of chlorophyll and carotenoid contents (Supplemental Fig. S4, B–G) (Arias et al., 2006). *35S::mHvPDS* also showed resistance to fluridone in the suppression of PR growth and induction of LR formation (**Fig. 1, C and D**). These results indicate that PDS is the target of fluridone for the suppression of PR growth and induction of LR formation. Subsequently, carotenoid-deficient mutants were examined for fluridone sensitivity. The phytoene synthase mutant *psy* showed reduced PR growth and LR formation (Van Norman et al., 2014) (Supplemental Fig. S1). PDS is encoded by a single gene, *PDS3*, in *Arabidopsis thaliana* (Qin et al., 2007), and its T-DNA insertion mutant, *pds3-1* (McElver et al., 2001; Meinke, 2019), also showed reduced PR growth rate and LR density (**Fig. 1, E– G**). The *pds3-1* plants were insensitive to fluridone for both PR growth suppression and LR density induction (**Fig. 1, E–G**), further indicating that PDS is a target for fluridone. Unlike the *pds3-1* plants, the *psy* plants showed weak resistance to fluridone compared to *pds3-1*, and their LR formation was induced by fluridone (Supplemental Fig. S1, A and B). These results indicate that the change in root architecture induced by fluridone is specifically dependent on the activity of PDS. Fluridone suppressed PR growth immediately after application, and the rate of suppression increased after 60 h of treatment (Supplemental Fig. S6).

**Figure 1.**
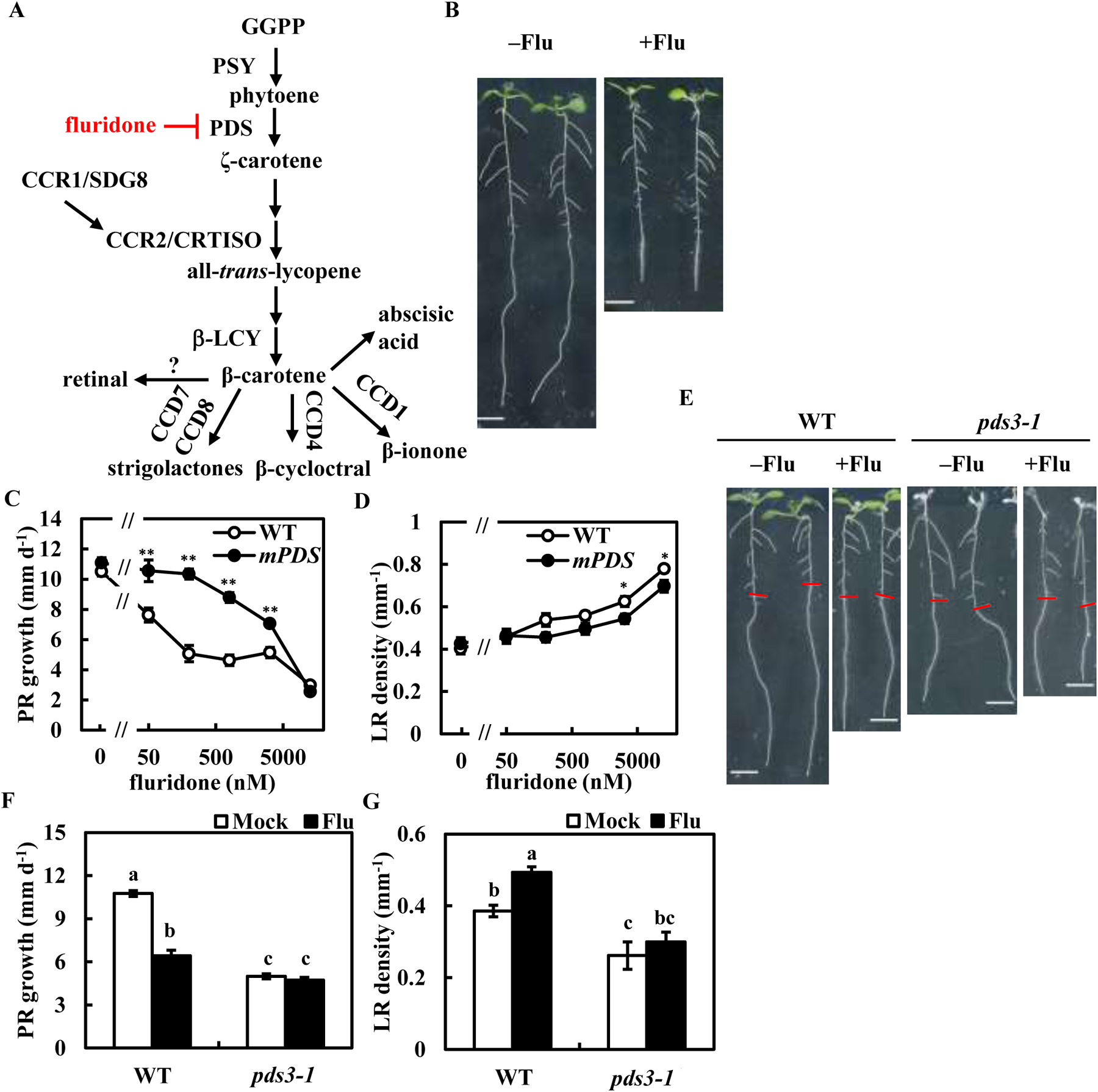
Fluridone inhibits phytoene desaturase activity to suppress PR growth. **A)** The carotenoid biosynthesis pathway. PSY, phytoene synthase; PDS, phytoene desaturase; CCR1/SDG8, carotenoid chloroplast regulatory1/set domain group 8; CCR2/CRTISO, carotenoid chloroplast regulatory2/crtiso; β-LCY, β-cyclase; CCD1/4/7/8, carotenoid cleavage dioxygenase 1/4/7/8. **B)** Five-day-old plants were transferred to medium with or without 12.8 µM fluridone. Photographs were captured at 3 d after transfer (dat). Scale bar = 5 mm. **C, D)** Five-day-old plants of WT and *mPDS* were transferred to medium with or without fluridone. **E–G)** Five-day-old plants of WT and *pds3-1* were transferred to medium with or without 800 nM fluridone. Photographs were captured at 3 dat. Scale bar = 5 mm. Red lines indicate the root tips at the time point of transfer. The PR growth rate in **(C, F)** was analyzed from 2 to 3 dat. The LR density in **(D, G)** was counted at 3 dat. Data represent the means ± standard error (SE) from 10-12 seedlings. *Significant differences compared to WT in **(C, D)** (*: *P* < 0.05, **: *P* < 0.01; Student’s *t-*test). Different lowercase letters above the bars in **(F, G)** indicate significant differences at *P* < 0.05 (one-way Analysis of Variance [ANOVA] following Tukey-Kramer test).

Fluridone inhibits the biosynthesis of ABA and SLs by inhibiting their precursor β-carotene (Yoshioka et al., 1998; López-Ráez et al., 2008). ABA and SLs promote root growth at low doses (Zhang et al., 2010; Ruyter-Spira et al., 2011). Therefore, we investigated whether exogenous ABA or the SL analog *racemic*-GR24 (*rac*-GR24) could complement fluridone-induced PR growth inhibition. The growth suppression of PR was not reversed by neither ABA nor *rac*-GR24 (Supplemental Fig. S5, G and H). The biosynthesis and signaling mutants of ABA or SLs were also tested for fluridone sensitivity. All the mutants examined showed comparable responses to fluridone compared to WT (Supplemental Fig. S5, A–C). These results suggest that ABA and SLs play a minor role in fluridone-induced PR growth suppression. The suppression of PR growth suggests an activation of auxin signaling by fluridone, as the increase in LR formation is under the control of auxin signaling (Du and Scheres, 2018).

### Fluridone increases endogenous auxin levels and activates auxin signaling

The levels of endogenous auxin in roots were measured during the temporary inhibition of PDS activity. The WT plants showed an increase in IAA levels after 8 h and 24 h of fluridone treatment (**Fig. 2**; Supplemental Fig. S7A). In contrast, the *35S::mHvPDS* plants showed no significant change of it (Supplemental Fig. S7B). The expression of the artificial auxin reporter *DR5::luciferase* (*DR5:LUC*) (Moreno-Risueno et al., 2010) was then measured. The *DR5:LUC* expression was transiently activated at the root tip regardless of the presence or absence of fluridone, which may be a response to plant stress (**Fig. 3**). After this transient activation, the *DR5:LUC* expression in fluridone-treated WT plants remained higher than in mock-treated plants after 4 h (**Fig. 3A**), while no such induction was detected in *35S::mHvPDS* plants (**Fig. 3B**). These results indicate that the fluridone-activated auxin response, like the induction of endogenous auxin, depends on PDS activity. The expression of *DR5:LUC* in fluridone-treated WT plants, but not in *35S::mHvPDS* plants, was reduced to below the mock level after 140 h (**Fig. 3**). This data suggests that the temporary inhibition of PDS activity leads to an increase in auxin levels and activation of the auxin signaling pathway, whereas long-term inhibition of PDS activity reduces the auxin response, possibly due to a carotenoid deficiency.

**Figure 2.**
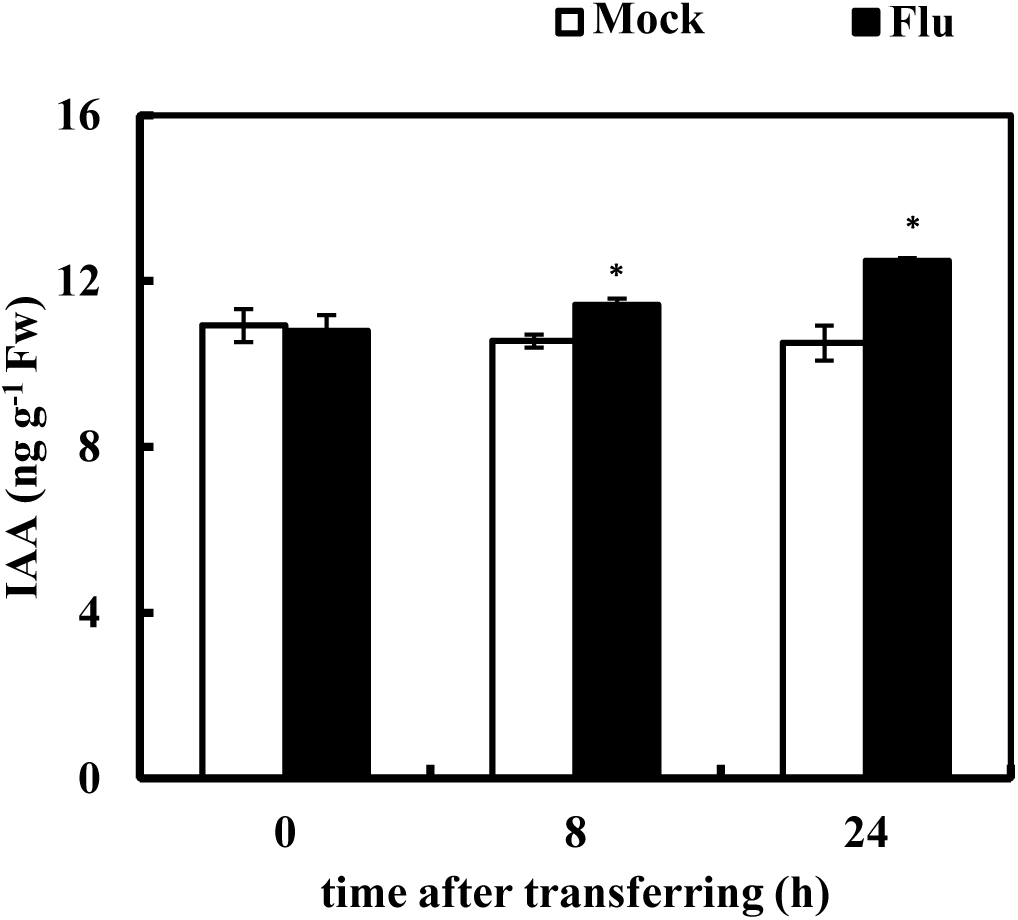
Fluridone induces endogenous IAA levels. Five-day-old plants were transferred to medium with or without 800 nM fluridone at time 0. The IAA contents in the roots were measured at the indicated time points. Data represent the means ± SE from 3 biological replicates. *Significant differences compared to mock-treated plants (*: *P* < 0.05; Student’s *t*-test).

**Figure 3.**
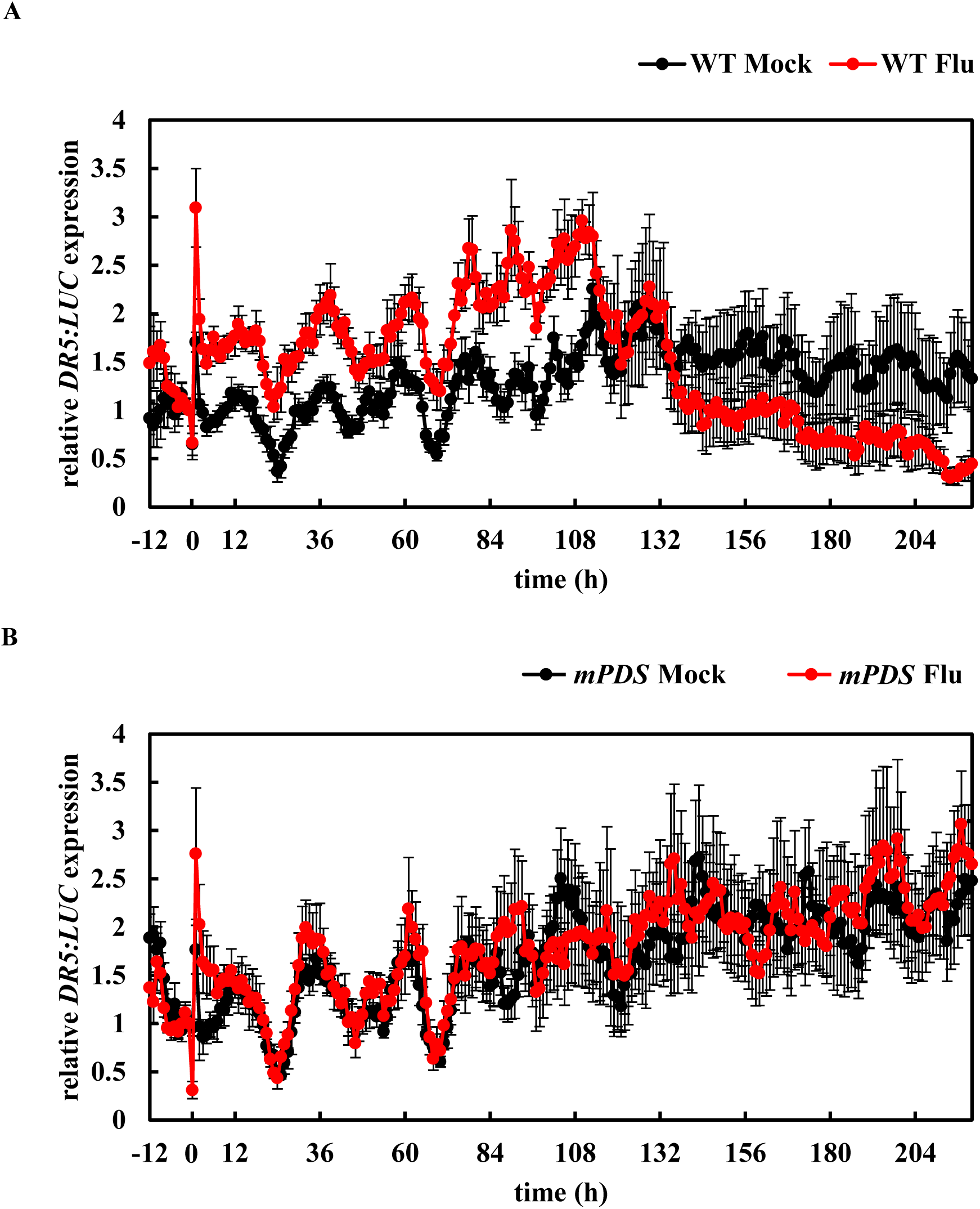
The expression of *DR5:LUC* at the root tips was increased following fluridone treatment. Four-day-old plants grown on 100 µM D-luciferin medium were transferred to D-luciferin-containing medium for a 1 d preincubation. Then plants were transferred to new D-luciferin-containing medium with or without 800 nM fluridone at time 0. The *DR5:LUC* expression in WT **(A)** and *mPDS* **(B)** was measured at an interval of 1 h. Data represent the means (relative to the mean of time –1 h) ± SE from 6 seedlings.

### Carotenoid-mediated regulation of root growth requires auxin biosynthesis and signaling

It is hypothesized that the temporary inhibition of PDS activity suppresses PR growth by increasing endogenous auxin levels and activating auxin signaling. To test this hypothesis, mutants in auxin signaling or auxin biosynthesis were examined. Indeed, the auxin permease mutant, *aux1-21*, and the ubiquitin-activating enzyme E1 mutant, *axr1-12* (Lincoln et al., 1990), were resistant to fluridone. *AUX/IAA* gain-of-function mutants *aux/iaa19/msg2-1* (Tatematsu et al., 2004) and *aux/iaa14/slr-1* (Fukaki et al., 2002), and the downstream transcription factor mutant *arf7-1 arf19-1* (Okushima et al., 2005) were weakly resistant to fluridone (**Fig. 4A**). *axr1-12* and *aux1-7* are reported to show a high level of auxin resistance on root elongation, whereas *msg2-1*, *slr-1*, and *arf7-1 arf19-1* exhibit relatively low or no resistance to auxin on root growth (Timpte et al., 1995; Fukaki et al., 2002; Tatematsu et al., 2004; Okushima et al., 2005). These results suggest that there is a correlation between fluridone resistance and auxin resistance in the auxin mutants.

**Figure 4.**
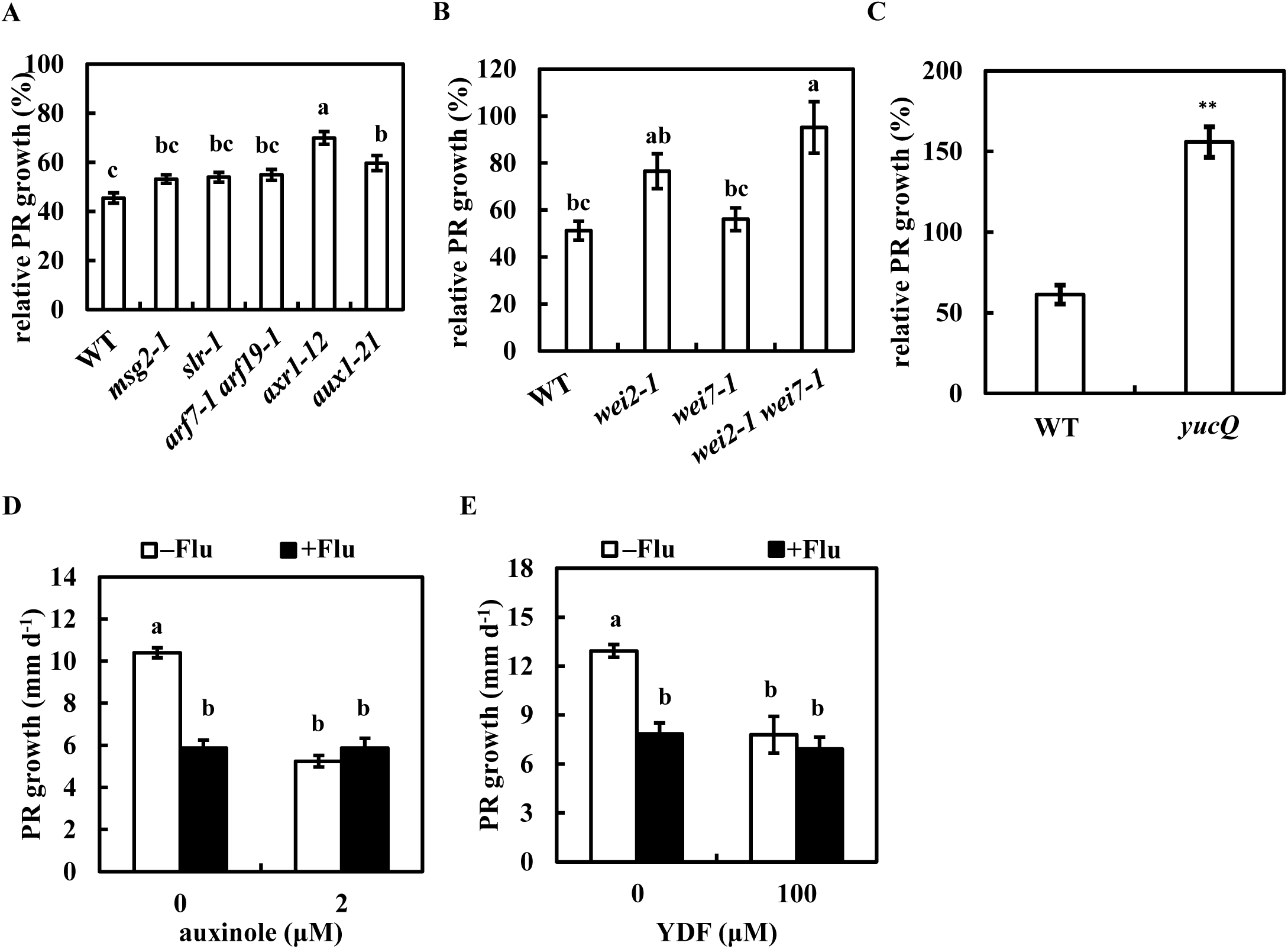
Auxin biosynthesis, signaling, and influx transport are required for fluridone-mediated root growth regulation. **A –C)** The resistance to fluridone in auxin-related mutants. Five-day-old plants were transferred to medium with or without 800 nM fluridone. **(A)** auxin signaling and influx transport mutants. **(B)** *wei2*, *wei7* and *wei2 wei7* mutants. **(C)** *yucQ* mutant. **D**, **E)** Five-day-old plants were transferred to medium with or without 800 nM fluridone and 2 µM auxinole **(D)** or 100 µM YDF **(E)**. The PR growth rate was analyzed from 2 to 3 dat. The relative PR growth rate in (A–C) was compared to mock-treated plants whose root growth rate is listed in Supplemental Table S1. Data represent the means ± SE from 20 seedlings in **(A)** and from 8 seedlings in **(B–E)**. Different lowercase letters above the bars in **(A**, **B, D, E)** indicate significant differences at *P* < 0.05 (one-way ANOVA following Tukey-Kramer test). *Significant differences compared to WT in **(C)** (**: *P* < 0.01; Student’s *t*-test).

Optimal levels of auxin are required for root growth, and both supraoptimal and suboptimal levels of auxin reduce root growth (Zhao, 2018). The root growth of the *wei2-1 wei7-1* double mutant was insensitive to fluridone, while its single mutant showed moderate or no resistance to fluridone, indicating the redundancy of these genes (**Fig. 4B**). The quintuple *YUCCA* mutant, *yucQ*, exhibited a severe root defect due to a reduction in endogenous auxin levels. Surprisingly, fluridone tended not to suppress the PR growth in the *yucQ* plants but did promote it (**Fig. 4C**; >100% of inhibition). This suggests that the inhibition of PDS activity increases endogenous auxin levels (**Fig. 2**), which compensates for the growth defect in the roots of *yucQ* (Chen et al., 2014). None of the *yuc* single mutants showed resistance to fluridone, indicating the redundancy of *YUCCA* genes (Supplemental Fig. S8, A and B). These results suggest that PDS activity negatively regulates endogenous auxin levels.

Pharmaceutical interference with auxin signaling and biosynthesis was then combined with the effect of fluridone. Auxinole is an antagonist of the auxin receptor TRANSPORT INHIBITOR RESPONSE 1/AUXIN SIGNALING F-BOX PROTEIN (TIR1/AFB) (Hayashi et al., 2012). Yucasin difluorinated analog (YDF) is a competitive inhibitor of YUCCA flavin monooxygenase (Tsugafune et al., 2017). Both auxinole and YDF suppressed PR growth, but the addition of fluridone to either inhibitor did not enhance the suppression of PR growth (**Fig. 4, D and E**). These results support the hypothesis that fluridone-induced suppression of PR growth requires an increase in endogenous auxin and proper auxin signaling. An inhibitor of auxin efflux transport, *N*-1-naphthylphthalamic acid (NPA), also suppressed PR growth in a dose-dependent manner (Reed et al., 1998). In contrast to auxinole and YDF, NPA showed an additive effect to that of fluridone in suppressing PR growth (Supplemental Fig. S9B), suggesting that the efflux transporter may not be the primary target of PDS inhibition.

### Negative control of carotenoid levels by auxin

The albino phenotype in the *pds3-1* and *psy* plants (**Fig. 1E**; Supplemental Fig. S1C) indicates the essential role of carotenoid biosynthesis in chloroplast development (Qin et al., 2007). The development of such chloroplasts is repressed by auxin signaling (Kobayashi et al., 2012). *arf7/msg1* and *axr1* are resistant to 2,4-D-induced chlorosis (Watahiki and Yamamoto, 1997), suggesting that carotenoid contents are also negatively regulated by auxin. Auxinole and YDF increased β-carotene levels in roots, whereas the application of fluridone abolished the β-carotene levels regardless of the presence of auxinole or YDF (**Fig. 5, A and B**). The auxin-deficient mutants, *wei2-1 wei7-1* and *yucQ* showed doubled β-carotene levels compared to WT (**Fig. 5C**), while auxin signaling mutants showed insignificant increased β-carotene levels (Supplemental Fig. S10C) which may be due to the redundant functions in these genes. These data suggest that endogenous auxin levels and signaling negatively control β-carotene levels. In addition, this negative regulation of carotenoid levels by auxin was also observed in the shoots (Supplemental Fig. S10, A and B).

**Figure 5.**
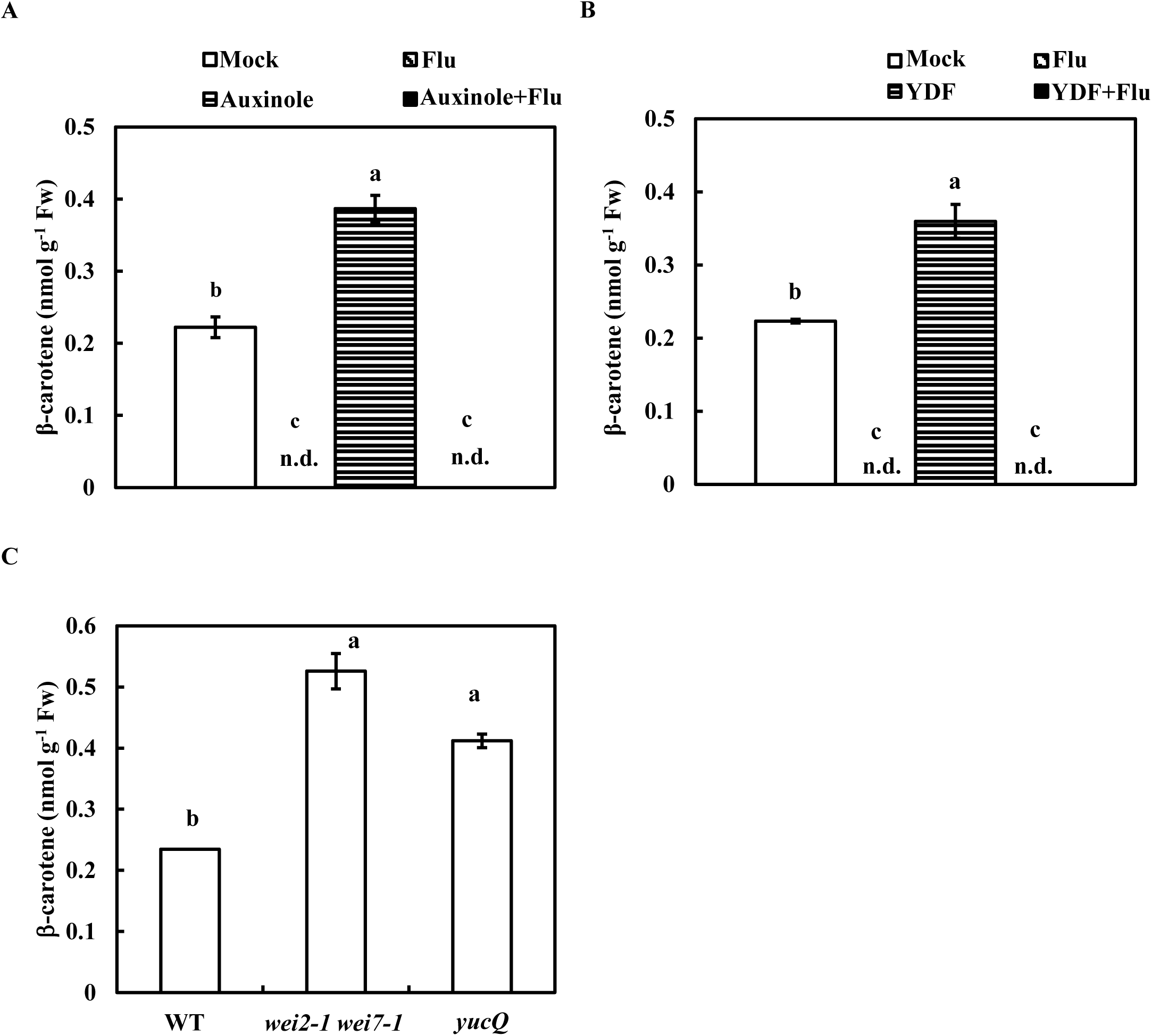
The carotenoid accumulation in the roots is negatively regulated by auxin levels and signaling. **A**, **B)** Five-day-old plants were transferred to medium with or without 800 nM fluridone and 10 µM auxinole **(A)** or 100 µM YDF **(B)**. The levels of β-carotene in the roots were measured at 2 dat. The bars indicate mock (white), fluridone (diagonal striped black), auxinole **(A)** or YDF **(B)** (horizontal striped black), and auxinole + fluridone **(A)** or YDF + fluridone **(B)** (black). **C)** The β-carotene levels of the auxin-deficient mutants in the roots were measured at 7 d after germination. Data represent the means ± SE from 3 biological replicates. Different lowercase letters above the bars in **(A**–**C)** indicate significant differences at *P* < 0.05 (one-way ANOVA following Tukey-Kramer test). n.d. indicates no detectable data.

### Expression of auxin- and carotenoid-related genes

To investigate whether PDS activity regulates auxin biosynthesis and signaling at the gene expression level, the expression of auxin-related genes in roots was measured after fluridone treatment. The expression of an early auxin-induced gene, *AUX/IAA19* or auxin biosynthesis gene, *YUCCA9*, increased approximately 4 and 2.5 times, respectively, after 4 h of fluridone treatment in WT, but not in *35S::mHvPDS* (**Fig. 6, A and B**). The induced expression of these genes was associated with the induction time of the *DR5:LUC* expression (**Fig. 3A**). These data suggest that the temporary inhibition of PDS activity activates auxin biosynthesis and signaling.

**Figure 6.**
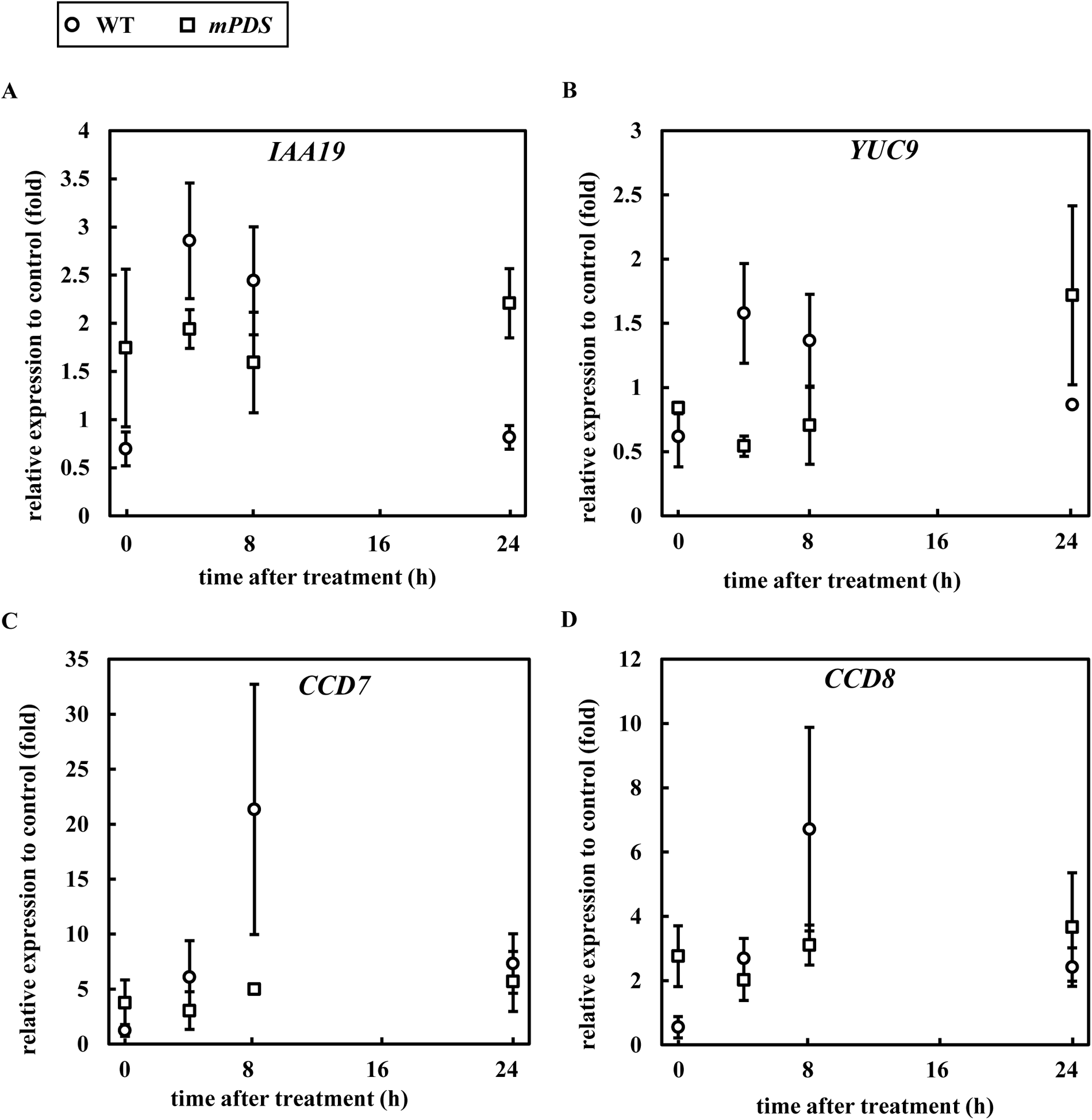
Fluridone activates auxin biosynthesis, response, and carotenoid catabolism genes. Five-day-old plants of WT and *mPDS* were transferred to medium with or without 800 nM fluridone at time 0. **A**–**D)** Relative expression of *IAA19*, *YUC9*, *CCD7*, and *CCD8* after fluridone treatment in the WT (open circle) and *mPDS* (open square). Data represent the means ± SE from 3 biological replicates.

The expression of genes in carotenoid biosynthesis (*CAROTENOID CHLOROPLAST REGULATOR2* [*CCR2*]), *CCR1* which regulates the expression of *CCR2* (Ruiz-Sola and Rodríguez-Concepción, 2012), and the carotenoid catabolism pathway (*CAROTENOID CLEAVAGE DIOXYGENASE 1* [*CCD1*], *CCD7* and *CCD8*) (Yuan et al., 2015) (**Fig. 1A**) was examined following fluridone treatment. The expression of *CCD7* and *CCD8* was strongly induced at 8 and 24 h in WT plants (**Fig. 6, C and D**). In contrast, the expression of *CCR1*, *CCR2* and *CCD1* showed little induction in comparison to *CCD7* and *CCD8* (Supplemental Fig. S11). These data suggest that after the temporary inhibition of PDS activity, carotenoid catabolism is activated rather than biosynthesis. *CCD7* and *CCD8* are reported to be induced by auxin (Bainbridge et al., 2005; Hayward et al., 2009). The GENEVESTIGATOR database (https://genevestigator.com) (Hruz et al., 2008) also shows that *CCD7* expression is induced within 0.5 to 1 h following exogenous IAA treatment (Supplemental Fig. S15). These suggest that the increased auxin levels by the inhibition of PDS activity likely feedback to upregulate carotenoid catabolism.

### Fluridone-mediated PR growth regulation in relation to retinal and auxin levels

Recent evidence has shown that the apocarotenoids β-cyclocitral and retinal, both derived from the oxidation of β-carotene, regulate root architecture (Olson and Hayaishi, 1965; Rodrigo et al., 2013; Dickinson et al., 2019; Dickinson et al., 2021). Fluridone potentially blocks the biosynthesis of such apocarotenoids and may result in the suppression of PR growth. Therefore, both β-cyclocitral and retinal were investigated to complement the suppression of PR growth by fluridone application. Retinal, but not β-cyclocitral, fully restored the PR growth suppression induced by fluridone (**Fig. 7A**; Supplemental Fig. S12). In addition, retinal also promoted the growth of PRs in the *pds3-1* mutant (**Fig. 7B**), suggesting that retinal is responsible for the short root phenotype of *pds3-1*. This also supports the hypothesis that the retinal is responsible for the suppression of root growth by fluridone. To further investigate the relationship between auxin signaling and retinal, we examined the expression of the auxin reporter *pIAA19::emerald luciferase-PEST* (*pIAA19::Eluc-PEST*) (Yamamoto et al., 2017), which was induced two-fold at the root tip by 10 nM 1-naphthalene acetic acid (NAA), a synthetic auxin analog. However, this induction was suppressed by the cotreatment with retinal (**Fig. 7E**). This result indicates that retinal suppresses the exogenous auxin response and raises the hypothesis that retinal may complement the inhibitory effect of auxin on root growth. Indeed, retinal restored the NAA-induced PR growth suppression in WT (**Fig. 7B**). Conversely, the short PR phenotype in *wei2-1 wei7-1* and *yucQ* was enhanced by the application of retinal (**Fig. 7C**). This suggests a further suppression of auxin levels by retinal in these mutants. In addition, retinal also showed an antagonistic effect to auxin on the LR formation induced by NAA and tryptophan (Supplemental Fig. S13, D and E). *SUPERROOT2/FEWER ROOT SUPPRESSOR1/CYP83B1* (*SUR2/FSP1*) is one of the cytochrome P450 gene family members of the indole-3-acetaldoxime (IAOx) pathway, which represents a non-IPyA pathway for IAA biosynthesis (Barlier et al., 2000; Sugawara et al., 2009). *sur2* mutant and the weaker allele *fsp1* mutant exhibit high LR density due to an overaccumulation of IAA (Delarue et al., 1998; Goto et al., 2023). Such a high LR density of *sur2* was also reduced by retinal (Supplemental Fig. S13G). These data are consistent with a negative regulation of endogenous auxin levels by retinal. However, no significant reduction in IAA levels was detected in the roots of retinal-treated WT plants (Supplemental Fig. S13I). Kakeimide (KKI), an inhibitor of the IAA-amino acid conjugating enzyme GRETCHEN HAGEN 3 (GH3s), accumulates endogenous IAA rapidly (Fukui et al., 2022) and suppressed PR growth in a dose-dependent manner (**Fig. 7D**), similar to the treatment with exogenous auxin (Supplemental Fig. S13C). However, unlike the case of exogenous auxin (**Fig. 7B**; Supplemental Fig. S13D), retinal did not counteract the KKI-induced suppression of PR growth and induction of LR formation (**Fig. 7D**; Supplemental Fig. S13H). Although the levels of retinal in fluridone-treated plants remain to be determined, these results suggest that retinal may target the degradation of auxin, thereby exerting an antagonistic effect on auxin.

**Figure 7.**
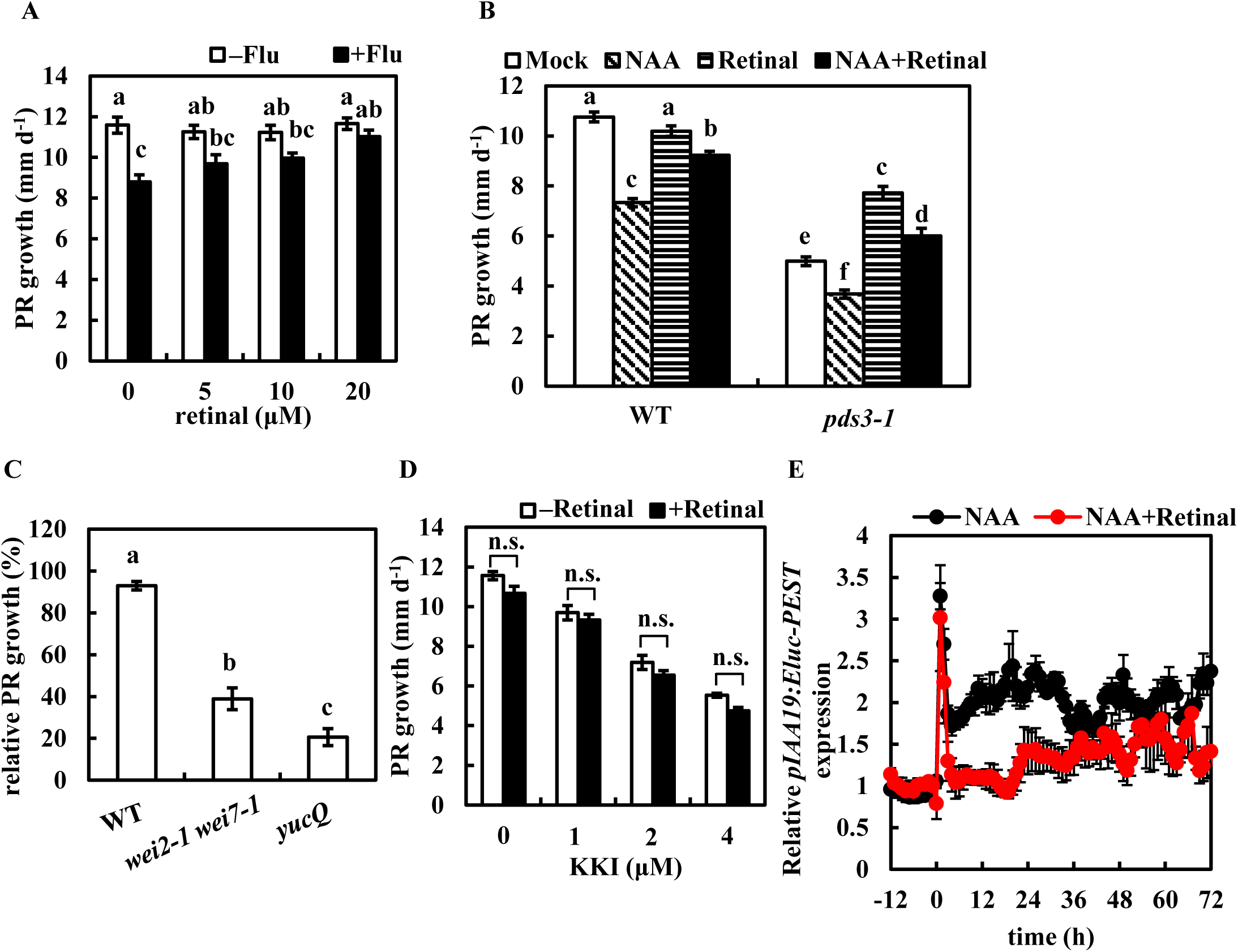
Retinal restores fluridone- and auxin-induced suppression of PR growth. **A)** Five-day-old plants were transferred to new medium with or without 800 nM fluridone and retinal. **B)** Five-day-old plants of WT and *pds3-1* were transferred to medium with or without 20 µM retinal and 20 nM NAA. **C)** Five-day-old plants of WT and the auxin-deficient mutants were transferred to medium with or without 20 µM retinal. **D)** Five-day-old plants were transferred to medium with or without 20 µM retinal and KKI. The PR growth rate was analyzed from 1 to 2 dat in **(A)** and 2 to 3 dat in **(B–D)**. The bars in **(B)** indicate mock (white), 20 nM NAA (diagonal stripe black), 20 μM retinal (horizontal striped black) and NAA + retinal (black). The relative growth in **(C)** was compared to mock-treated plants. **E)** The relative *pIAA19::Eluc-PEST* expression by cotreatment of NAA and retinal. Four-day-old plants grown on 100 µM D-luciferin medium were transferred to new D-luciferin-containing medium for a 1 d preincubation. Then plants were transferred to new medium containing 100 µM D-luciferin, 10 nM NAA and with or without 20 µM retinal at time 0. The *pIAA19* expression was measured at an interval of 1 h. Data represent the means ± SE from 8-10 seedlings in **(A**–**D)** and the means (relative to the means of time –1 h) ± SE from 6 seedlings in **(E)**. Different lowercase letters above the bars in **(A–C)** indicate significant differences at *P* < 0.05 (one-way ANOVA following Tukey-Kramer test).

## Discussion

### Inhibition of PDS activity increases the levels of auxin in roots

The regulation of root growth by temporary inhibition of carotenoid biosynthesis was the subject of this study. Here we showed that a temporary application of fluridone increased endogenous auxin levels in the roots but not in the shoots (**Fig. 2**; Supplemental Table 2). Such a temporary inhibition of PDS activity may exceed the optimal concentration of auxin levels for the root growth, thereby resulting in the suppression of PR growth and the promotion of LR formation (**Fig. 1, C and D**). The endogenous auxin levels in *yucQ* were suboptimal, and the short roots of *yucQ* could be rescued by a low dose application of exogenous IAA (5 nM) (Chen et al., 2014). The partial recovery of root growth in the short roots of *yucQ* correlates with the root-specific increase in auxin levels by the inhibition of PDS activity (**Fig. 2 and 4C**). The *sur2/fsp1* and *yuc1D* plants accumulate endogenous auxin and increase the growth of hypocotyls (Delarue et al., 1998; Zhao et al., 2001). However, the length of hypocotyls was similar between fluridone-treated and mock-treated plants (data not shown), confirming a root-specific induction of auxin by fluridone. Like the *yucQ* plants, the *wei2 wei7* (*asa1 asb1*) plants showed a reduction in endogenous IAA levels and a short root phenotype. Although the root of *wei2 wei7* showed resistance to fluridone, it did not restore the root growth of *wei2-1 wei7-1*, in contrast to that of *yucQ* (**Fig. 4, B and C**). A low dose application of anthranilic acid, a precursor of IAA and tryptophan, partially recovered the root growth in *wei2-1 wei7-1* without rescuing its endogenous IAA levels (Doyle et al., 2019). This suggests that the short roots of the *wei2-1 wei7-1* plants require anthranilic acid rather than increasing the suboptimal concentration of IAA to compensate their growth. In fact, a higher concentration of anthranilic acid significantly but not completely rescued root growth of *wei2-1 wei7-1* (Doyle et al., 2019). Low dose application of exogenous NAA did not rescue or inhibit root growth in *wei2-1 wei7-1* (Supplemental Fig. S8C), further supporting that endogenous auxin is suboptimal and it is not a primary requirement for rescuing the PR growth in *wei2-1 wei7-1*. The inhibition of auxin signaling and biosynthesis by auxinole and YDF, respectively, suppressed PR growth; however, fluridone addition did not further enhance this suppression (**Fig. 4, D and E**). In contrast to auxinole and YDF, exogenous auxin and fluridone exhibited an additive effect on PR growth suppression (Supplemental Fig. S9A). This suggests that the optimal level of auxin and auxin signaling are required for PR growth, and that excessive or insufficient levels of auxin can reduce PR growth. Taken together, fluridone-induced PDS inhibition increases root auxin levels, activates auxin signaling, and thus suppresses PR growth.

### The roles of apocarotenoids on fluridone-mediated root growth

The temporary inhibition of PDS activity triggers the auxin-mediated suppression of PR growth. However, the downstream product of PDS activity responsible for this process has yet to be identified. The extremely hydrophobic nature of β-carotene made it impossible for the complementation test by exogenous application to the plants (data not shown), which is inhibited by fluridone. Although ABA and SLs are the candidates for their close interactions with auxin in the regulation of root architecture (Ruyter-Spira et al., 2011; Emenecker and Strader, 2020), this study provides no evidence that they are responsible for the temporary inhibition of PDS activity on the root architecture regulation (Supplemental Fig. S5). Consistently, the LR formation of the excised tomato roots is also induced by fluridone, even in the presence of ABA (Hooker and Thorpe, 1998).

Dickinson et al. (2021) report that retinal (1 μM) induces LR capacity. In this study, the application of a higher dose of retinal (20 μM) showed antagonistic effects to auxin on the PR growth and the LR formation (**Fig. 7, B and E**; Supplemental Fig. S13, C, D, E and G). Although retinal partially complemented the short roots of the *pds3-1* plants (**Fig. 7B**), it did not alter LR density (Supplemental Fig. S13D). 20 μM retinal also did not alter PR growth or LR density in the WT plants (**Fig. 7B**; Supplemental Fig. S13D). Instead, the sites of LR emergence in retinal-treated WT plants were shifted away from the root tip compared to those of mock-treated WT plants (Supplemental Fig. S13F). Therefore, it is speculated that retinal suppressed the auxin response during the development of the LR primordia and delayed the emergence of LRs, resulting in longer zones of unbranched PR. It should be noted that Dickinson et al. (2021) measure the LR capacity by excising the root tips to encourage LR formation which evokes the root cut response (Xu et al., 2017), which differs from this study. The severe short root phenotype in *yucQ* is further repressed by YDF due to a further inhibition of auxin biosynthesis in *yucQ* (Tsugafune et al., 2017). Although retinal did not consistently alter endogenous free IAA levels (Supplemental Fig. S13I), it further suppressed PR growth in the auxin-deficient mutants (**Fig. 7C**) and had an additive effect on the YDF-induced suppression of root growth (data not shown). Instead, the antagonistic effect of retinal on auxin was abolished by the auxin catabolism inhibitor KKI (**Fig. 7D**; Supplemental Fig. S13H), suggesting that retinal may target downstream substrates of GH3s in the auxin catabolic pathway. The study of mutants in the auxin catabolic pathway on fluridone and retinal may be required for this confirmation. Although the expression of *YUC9* was enhanced by fluridone treatment (**Fig. 6B**), a single *yuc9* mutant did not show resistance to fluridone (Supplemental Fig. S8A). This suggests that the fluridone-induced accumulation of endogenous IAA may be primarily contributed by the suppression of the auxin catabolic pathway rather than by the activation of the auxin biosynthesis pathway.

### The difference between continuous carotenoid deficiency and temporary inhibition of PDS activity

Van Norman et al. (2014) and this study showed that both the *psy* and *pds3-1* plants had shorter PR lengths compared to WT (**Fig. 1, E and F**; Supplemental Fig. S1, A and C). Similarly, temporary exposure to fluridone reduced PR growth. Short-term fluridone treatment led to rapid and modest suppression of root growth, whereas long-term exposure (>60 h) resulted in accelerated suppression of root growth (Supplemental Fig. S6), which may correspond to the short PRs in *psy* and *pds3-1*. The root growth of fluridone-treated WT plants was fully restored by retinal on the second day after treatment, but the extent of root growth recovery in *pds3-1* was less than that of WT (Supplemental Fig. S14). By the third day after treatment (>60 h), such retinal-mediated recovery of root growth became ineffective in WT, whereas it remained at the same level in *pds3-1* (Supplemental Fig. S13A and S14). Temporary application of fluridone resulted in the degradation of chlorophylls, leading to chlorosis in established cotyledons and bleaching in newly emerged leaves (Supplemental Fig. S4A) (Kim et al., 2004), while the albino *pds3* mutant lost the ability to differentiate plastids (Qin et al., 2007). The difference in root growth recovery between the fluridone-exposed plants and the *pds3-1* plants may reflect the process of plastid degradation by fluridone and undifferentiated proplastids, respectively, suggesting a further role of proplastid differentiation for root growth.

Although both *psy* and *pds3-1* reduced the number of LRs (**Fig. 1G**; Supplemental Fig. S1B), fluridone increased the number of LRs in the pre-grown areas of PRs and decreased it in the newly grown areas of PRs in WT (Supplemental Fig. S3), which corresponds to the activation of the *DR5:LUC* expression in the upper part of PRs (Supplemental Movie. S1). An induction of LR formation by fluridone has been reported in the excised tomato roots (Hooker and Thorpe, 1998). It is speculated that the process of plastid degradation increases auxin levels in pre-grown roots, while newly grown roots remain undifferentiated proplastids and reduce LRs, as in the *psy* and *pds3-1* plants (Supplemental Fig. S3) (Van Norman et al., 2014). This hypothesis is consistent with the result that the *DR5:LUC* expression at the root tips was increased upon fluridone application, and eventually decreased upon long-term incubation (**Fig. 3A**).

### Conclusion

In this study, we found that the apocarotenoid retinal controls the growth of PRs through endogenous auxin levels (**Fig. 8**). Temporary inhibition of PDS activity blocks retinal biosynthesis and then leads to auxin accumulation to suppress root growth. Long-term exposure to fluridone further suppresses root growth, the underlying mechanisms of which remain to be elucidated. The role of retinal here may be different from that of promoting LR capacity (Dickinson et al., 2021). Retinal may regulate auxin catabolism and auxin homeostasis. In addition, auxin may further regulate carotenoid accumulation through a feedback mechanism involving the expression of genes associated with carotenoid catabolism. This model provides insight into the role of the apocarotenoid retinal in root growth regulation. The quantitative analysis of retinal and its derivatives, as well as their targets that may regulate auxin homeostasis, is warranted in the future.

**Figure 8.**
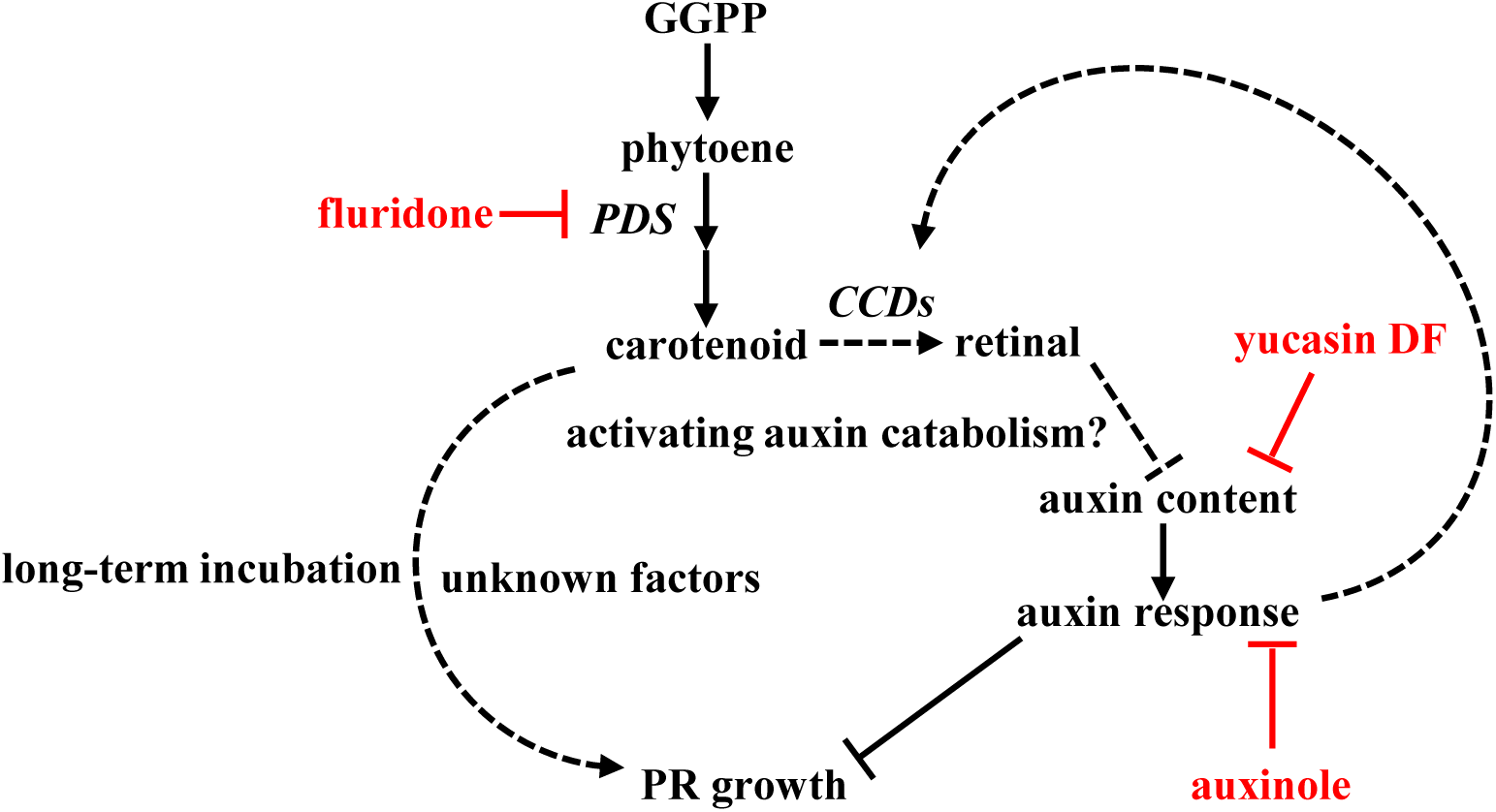
Putative model of carotenoid-mediated PR growth regulation. Under normal conditions, PR growth is controlled by auxin homeostasis. The short-term fluridone treatment inhibits PDS activity and blocks consequent retinal biosynthesis to increase endogenous auxin levels. The increase of auxin levels may exceed its optimal levels for root growth. Retinal may trigger the IAA catabolism pathway to control endogenous auxin levels. Under long-term fluridone incubation, the carotenoid deficiency leads to an acceleration of root growth suppression via unknown factors. The activated auxin response feedback for the carotenoid catabolism gene *CCD* activity in the regulation of carotenoid content. Black dashed lines indicate unknown or predicted mechanisms.

## Materials and methods

### Plant materials and growth conditions

The *Arabidopsis thaliana* mutants and WT plants as well as *PDS* transgenic plants were in the Columbia background, except for *yuc5*, which is in the Landsberg *erecta* background. *pds3-1* used in this work refers to *pigment defective 226* (*pde226-1*). The *arf7-1 arf19-1*, *slr-1*, *yuc1*, *yuc2*, *yuc3*, *yuc4-1*, *yuc5*, *yuc6*, *yuc7*, *yuc8*, *yuc9*, *yuc10*, and *yuc11* plants were described previously (Xu et al., 2017). The *axr1-12* (Lincoln et al., 1990), *aux1-21* (Roman et al., 1995), *aba1-6* (Niyogi et al., 1998), *aba2-1* (González-Guzmán et al., 2002), *psy* (Alonso et al., 2003) and *pde226-1* (McElver et al., 2001) plants were obtained from the Arabidopsis Biological Resource Center. *wei2-1*, *wei7-1*, *wei2-1 wei7-1* (Stepanova et al., 2005) and *sur2* (Delarue et al., 1998) plants were provided from Dr Hidehiro Fukaki (Kobe University). *aba3-2* (Koornneef et al., 1982), *abi1c* (Koornneef et al., 1984), *abi3-9*, *abi4-11*, *abi5-7* (Nambara et al., 2002), *ost1-1* (Mustilli et al., 2002) and *pyrQ* (Park et al., 2009) plants were provided from Dr Masanori Okamoto (RIKEN Center for Sustainable Resource Science). *max1-1*, *max2-1* (Stirnberg et al., 2002) and *max3-9* (Booker et al., 2004) plants were provided from Dr Kiyoshi Tatematsu (National Institute for Basic Biology). The *yucQ* mutant was a generous gift from Dr Hiroyuki Kasahara (Tokyo University of Agriculture and Technology) with an agreement from Dr Yunde Zhao (University of California San Diego). *DR5:LUC* (Moreno-Risueno et al., 2010) was kindly provided from Dr Philip N. Benfey (Duke University). In this study, *DR5:LUC* reporter in the *DR5:LUC/VENUS* line (Toyokura et al., 2019) was observed. Fluridone, norflurazon, deflufenican, picolinafen, and retinal were purchased from Fujifilm Wako Pure Chemical. KKI, YDF, and auxinole were provided by Kei-ichiro Hayashi. Seeds were surface-sterilized using chlorine gas for at least 45 min, suspended in 0.38% agarose, and sown on half-strength Murashige and Skoog (MS) medium (Duchefa Biochemie) supplemented with 1% (w/v) sucrose, 0.6% (w/v) gellan gum, and 0.5 mM MES (pH 5.8). Stratification was performed at 4°C for 3 d in the dark. Plants were grown on vertically oriented plates at 22°C under constant light if not specifically indicated (80–110 µmol m^−2^ s^−1^). Stock solutions of phytohormones and inhibitors were prepared in dimethyl sulfoxide and filtered through a 0.45 μM disc filter.

### PDS-resistant vector and plant transformation and screening

The plasmid pPDSer1303 was kindly provided by Renée S. Arias (USDA ARS, USA). The *pds* mutation vector construction was described previously (Arias et al., 2006). The plasmids were transformed into *Agrobacterium* strain C58C1 and used to transform *Arabidopsis thaliana* via the dip floral method. *Agrobacterium* harboring the plasmid pCambia1303 without *pds* was used as a control for the transformations and later as a control for herbicide screening. Seeds from T1, T2, and T3 generations were screened with hygromycin (50 mg/L) and fluridone (25 nM) separately. Seedlings that were resistant to hygromycin in T1 were selected in either hygromycin or fluridone (25 nM) screening in T2 and T3. T3 seeds were screened for resistance to fluridone at 25, 50, and 200 nM.

### Quantification of LR density and PR growth

Plants were grown on vertical plates for 5 d and then transferred to a new half-strength MS medium with or without chemicals indicated. Plant images were captured using a flatbed scanner (GT-X980 and DS-50000, EPSON) within 4 d after transfer. The LR density (the number of emerged LRs besides an adventitious root/the root length from the root–shoot junction to the youngest LR toward the root tip) (Dubrovsky and Forde, 2012) and the PR growth rate were analyzed with Fiji software (https://fiji.sc/).

### Quantification of PR growth via time-lapse imaging

Plants were grown on vertical plates for 4 d and then transferred to a new half-strength MS medium. Plant images were captured using a digital camera (Lumix G5, Panasonic). The interval was 30 min. After 1 d of photography, the seedlings were transferred to a new half-strength MS medium with or without 800 nM fluridone. The PR growth rate was analyzed using Fiji software (https://fiji.sc/).

### Quantification of IAA

Seeds were sown in sterilized 300 µm nylon mesh and transferred to control or fluridone plates after 5-d germination together with mesh. Shoot and root samples were harvested at specific time points after transfer and frozen in liquid nitrogen. Hormonome analysis was conducted as previously described (Xu et al., 2017). Briefly, samples of ∼100 mg fresh weight (Fw) were suspended in 80% (v/v) aqueous methanol with [^2^H_5_]-IAA as internal standards. The samples were homogenized and the supernatant was loaded onto a Bond Elut C18 cartridge (100 mg, 3 ml; Agilent Technologies) and eluted with 80% (v/v) aqueous methanol. The concentrated samples were subjected to liquid chromatography with electrospray ionization tandem mass spectrometry comprising a quadrupole tandem mass spectrometer (Agilent 6460 Triple Quadrupole mass spectrometer) with an electrospray ion source and an Agilent 1200 separation module. The raw data were extracted from MassHunter software (Agilent Technologies) and examined in Excel (Microsoft).

### Bioluminescence imaging analysis

D-Luciferin was purchased from Gold Biotechnology. Four-day-old plants grown on 100 µM D-luciferin medium were transferred to new D-luciferin medium for a 1 d preincubation and transferred to new D-luciferin medium with or without chemicals indicated. Time-lapse imaging was performed using a camera (ASI294MMPro, ZWO) with control software (SharpCap, v4.0). The images were captured at an interval of 1 h, including 15 min of exposure and 45 min of red and blue light (100 µmol m^−2^ s^−1^) and then saved in a TIFF format for further analysis using Fiji software (https://fiji.sc/).

### RNA isolation and quantitative reverse transcription polymerase chain reaction analysis

Seeds were sown on control medium with 300 μm nylon mesh. Roots were placed on plates containing no herbicide (mock) or 800 nM fluridone, root samples were harvested at 0, 4, 8, and 24 h and frozen in liquid nitrogen. Total RNA was extracted and purified using an RNA isolation kit (ISOSPIN Plant RNA, NIPPON GENE). Complementary DNA was synthesized from the total RNA according to the manufacturer’s instructions (ReverTra Ace qPCR RT Master Mix, Toyobo). Quantitative reverse transcription polymerase chain reaction (qRT– PCR) was performed in optical 96-well plates via the LightCycler 480 II system (Roche Life Science) using KOD SYBR qPCR Mix (Toyobo). Primer pairs spanning exon–exon junctions were designed using the QuantPrime program to avoid genomic DNA amplification as listed in Supplemental Table S3 (Xu et al., 2017). The specificity of the reactions was verified via melt curve analysis and capillary electrophoresis (Multina, Shimadzu). Standard curve analysis was used to evaluate the efficiency of the reactions. ACTIN2 was used as an internal standard. The qRT–PCR program included one cycle at 98℃ for 2 min, followed by 40 cycles of 98℃ for 10 s, 60℃ for 10 s, and 68℃ for 30 s. The cycle time values were determined using the second derivative maximum method (Tichopad et al., 2003) via LightCycler software (version 1.5, Roche Life Science). The data were analyzed using the comparative threshold cycle (Ct) method (2^−ΔΔCt^) (Schmittgen and Livak, 2008).

### Quantification of carotenoid via high-performance liquid chromatography (HPLC) analysis

Seeds were sown on control medium with 300 μm nylon mesh. The seedlings were grown under the conditions described above. Five days after germination, the seedlings together with nylon mesh were transferred to plates with or without chemicals indicated. Shoot tissues (∼20 mg) or root tissues (∼100 mg) were harvested at specific time points and stored at −80°C. Total carotenoids were extracted using acetone wako 1^st^ grade (FUJIFILM Wako Chemicals). Acetone was added to the tissue samples that had been mixed with beads and silica gel and shaken for 2 min. Total carotenoids were separated via centrifugation for 10 min at 14000 rpm at 4℃. High-performance liquid chromatography analysis for carotenoid quantification was performed as previously described (Zapata et al., 2000). An aliquot of the sample was subjected to a C8 column (4.6 × 150 mm, Waters Symmetry C8, Waters) equilibrated with A solution (methanol:acetonitrile:aqueous pyridine solution [0.25 M pyridine] = 2:1:1 [v:v:v]). Pigments were eluted with a linear gradient of 0%–40% B solution (methanol:acetonitrile:acetone = 1:3:1 [v:v:v]) for 2 min, 40%–95% for 6 min, an isocratic elution for 5 min, a linear gradient of 95%–0% for 2 min, and an isocratic elution for 7 min at a flow rate of 1 mL/min. Pigments were identified based on their retention time relative to known standards and quantified by peak area on the chromatogram.

### Statistical analysis

All data shown in graphs and tables are represented as the means ± SE from 3 replicates, except for Supplemental Fig. S2, which is performed once. The statistical significance is determined by 2-tailed Student’s *t*-test (*: *P* < 0.05, **: *P* < 0.01) or one-way ANOVA following Tukey-Kramer test (Different lowercase letters indicate significant differences, *P* < 0.05).

## Author contributions

Author contributions: K.X. conceived the project and designed the experiments, which were mostly carried out and analyzed by K.X.; H.Z. and F.L. supported retinal-related experiment design, process and data analysis; E.Y. and M.A. were responsible for IAA quantification and data collection; K.X. and M.K.W. wrote the article. K.X., H.Z., F.L., E.Y., M.A., K.H., H.F., H.I. and M.K.W. read, commented and approved the final manuscript.

## Acknowledgments

We thank Dr. Renée Arias (United States Department of Agriculture) and Dr. Franck Dayan (Colorado State University) for the supply of plasmid *35S::mHvPDS* and for the kind reviewing of the manuscript. This research was funded by grants-in-aid for scientific research from the Japan Society for the Promotion of Science (JSPS) (JP19H05673 to H. F. and M. K. W., JP21K19097 to K.H.), Japan Science and Technology Agency Support for Pioneering Research Initiated by the Next Generation (JST SPRING) (Grant Number JPMJSP2119 to K. Y.) and ACRO Research grant of Teikyo University (TeTe20-01 to M.A.).

