## Supplemental Figures 1-15 for "The exogenous application of the apocarotenoid retinaldehyde negatively regulates auxin-mediated root growth"

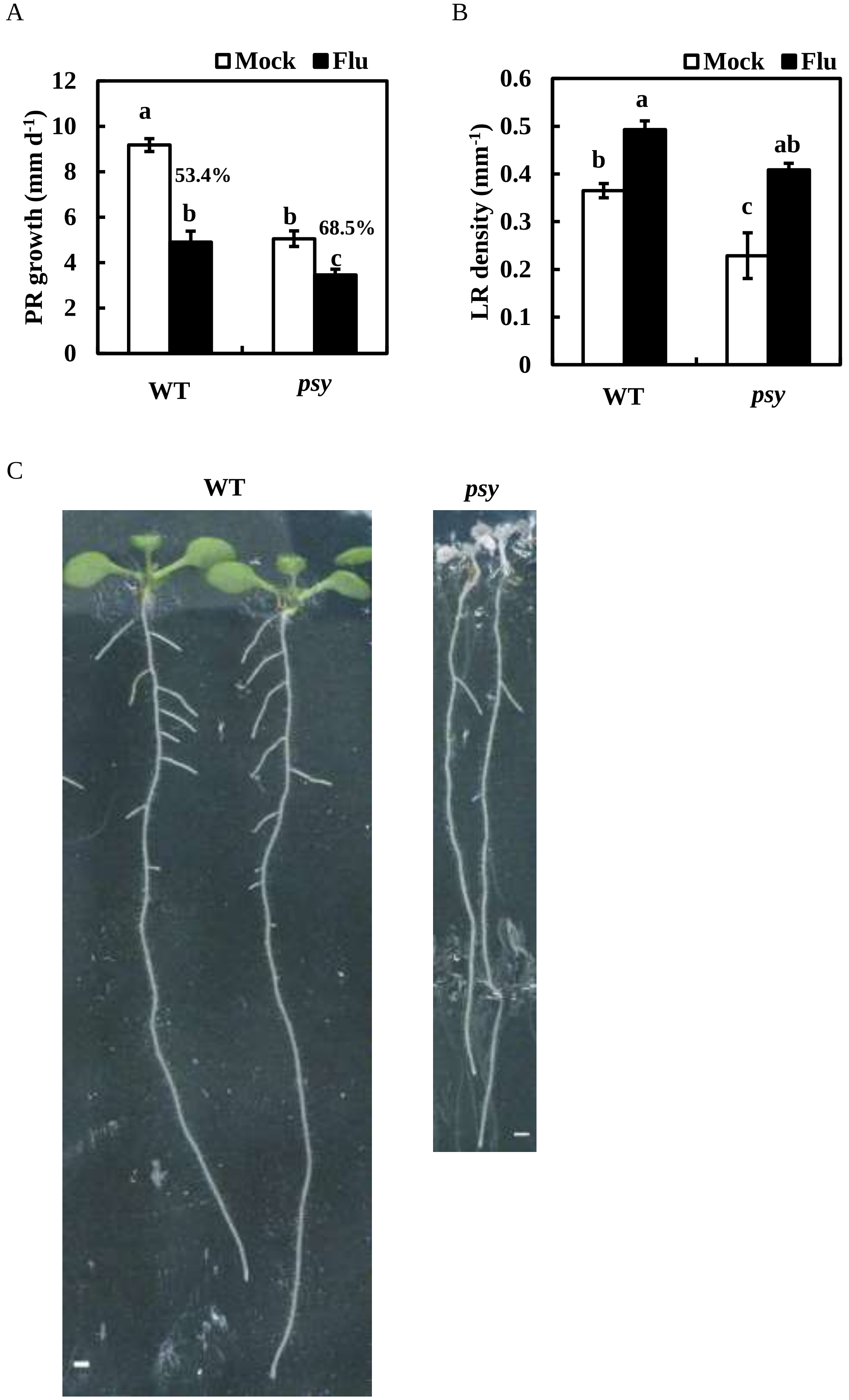


Supplemental Figure S1. The phenotypes and fluridone resistance of the *psy* mutant. A, B) Five-day-old plants of WT and *psy* were transferred to medium with or without 800 nM fluridone. The PR growth rate (A) was analyzed from 2 to 3 dat. The LR density (B) was counted at 3 dat. C) The pictures of WT and *psy* mutant captured at 8 d after germination (without transfer). Scale bar = 1 mm. The percentage in (A) means the relative PR growth rate compared to mock-treated plants. Data represent the means ± SE from 10 seedlings in (A, B). Different lowercase letters above the bars indicate significant differences at $P < 0.05$ (one-way ANOVA following Tukey-Kramer test).

A

B

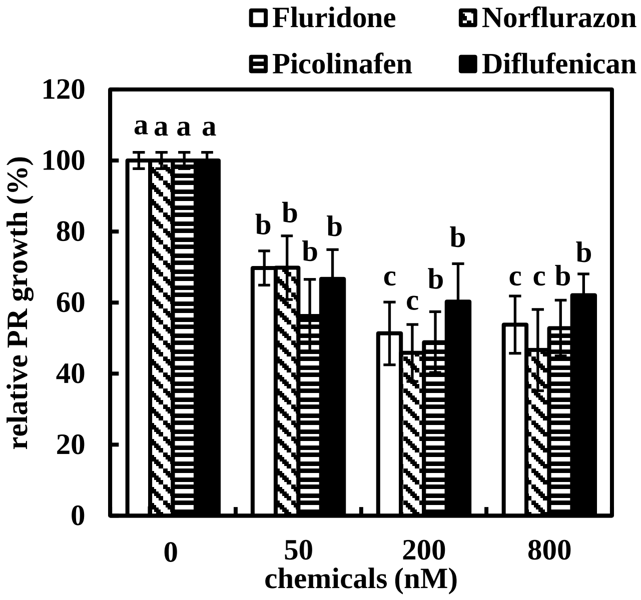


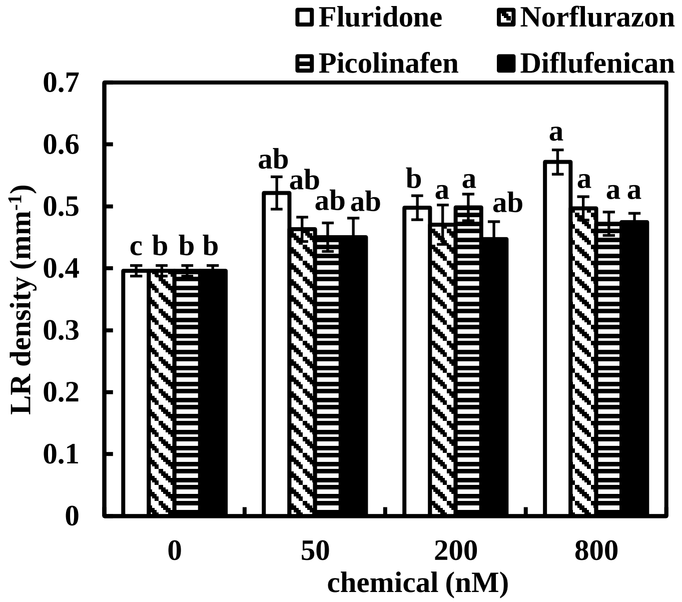


Supplemental Figure S2. PDS inhibitors exhibit a fluridone-like effect on PR suppression and LR induction. A, B) Five-day-old plants were transferred to medium with or without inhibitors. The relative PR growth rate compared to mock-treated plants (A) was analyzed from 2 to 3 dat. The LR density (B) was counted at 3 dat. The bars indicate fluridone (white), norflurazon (diagonal striped black), picolinafen (horizontal striped black), and diflufenican (black). Data represent the means ± SE from 8 to 14 seedlings. Different lowercase letters above the bars indicate significant differences at $P < 0.05$ (one-way ANOVA following Tukey-Kramer test).

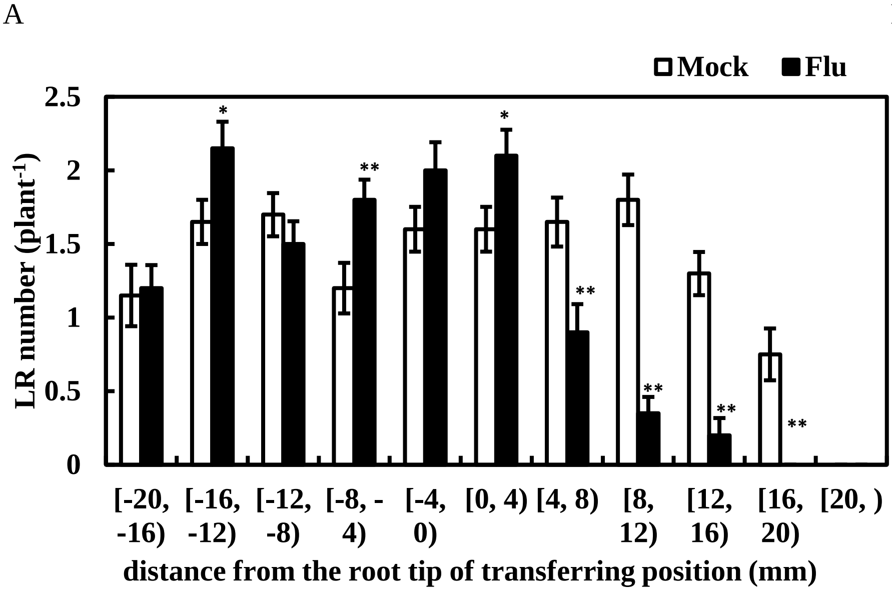


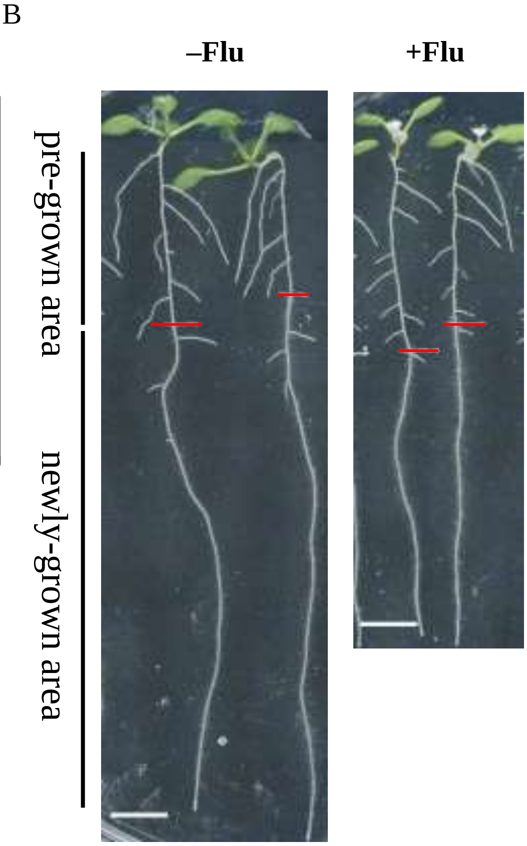


Supplemental Figure S3. Fluridone induces LR formation in the pre-grown areas of PR but severely reduces it in the newly grown areas of PR. A, B) Five-day-old plants were transferred to medium with or without 800 nM fluridone. The number of LRs was counted at 4 dat. The root tips of the transferring positions were positioned as 0. The pre-grown root was positioned as < 0. The newly grown root was positioned as > 0. Photographs were captured at 4 dat. Scale bar = 5 mm. Red lines indicate the root tips of the transferring position. Data represent the means ± SE from 20 seedlings. *Significant differences compared to mock-treated plants (*: $P < 0.05$, **: $P < 0.01$; Student's *t*-test).

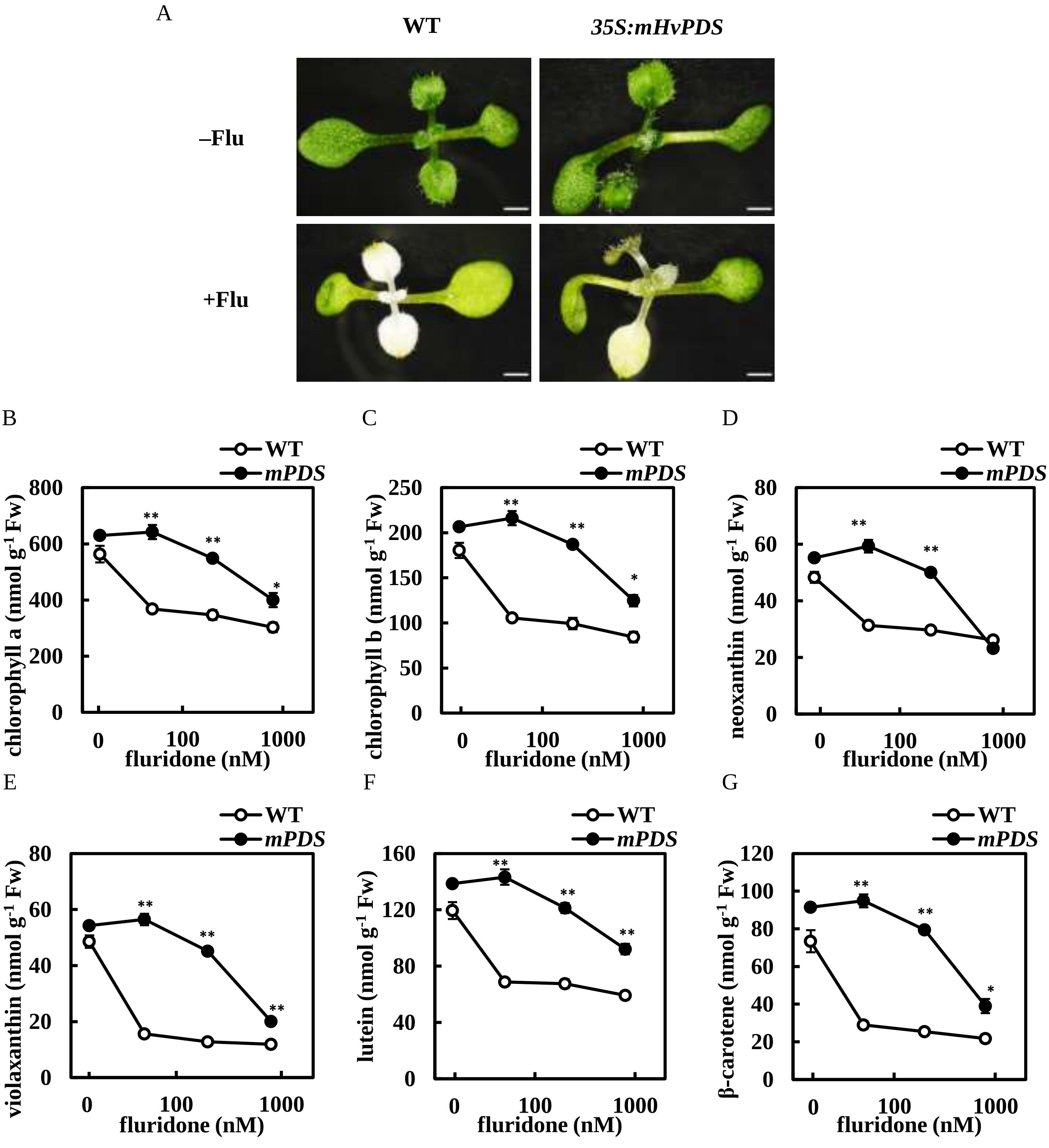


Supplemental Figure S4. Fluridone reduces chlorophyll and carotenoid contents, and *mPDS* plants are resistant to it. A–G) Five-day-old plants of WT and *mPDS* were transferred to medium with or without 800 nM fluridone. (A) Leaf pigment bleaching effect of WT and *mPDS* by fluridone. Photographs were captured at 3 dat. Scale bar = 5 mm. Chlorophyll a (B), chlorophyll b (C), neoxanthin (D), violaxanthin (E), lutein (F) and β-carotene (G) contents in the whole plants of WT (hollow circle) and *mPDS* (solid circle) were quantified at 2 dat. Data represent the means ± SE from 3 biological replicates. *Significant differences compared to WT (*: $P < 0.05$, **: $P < 0.01$; Student’s *t*-test).

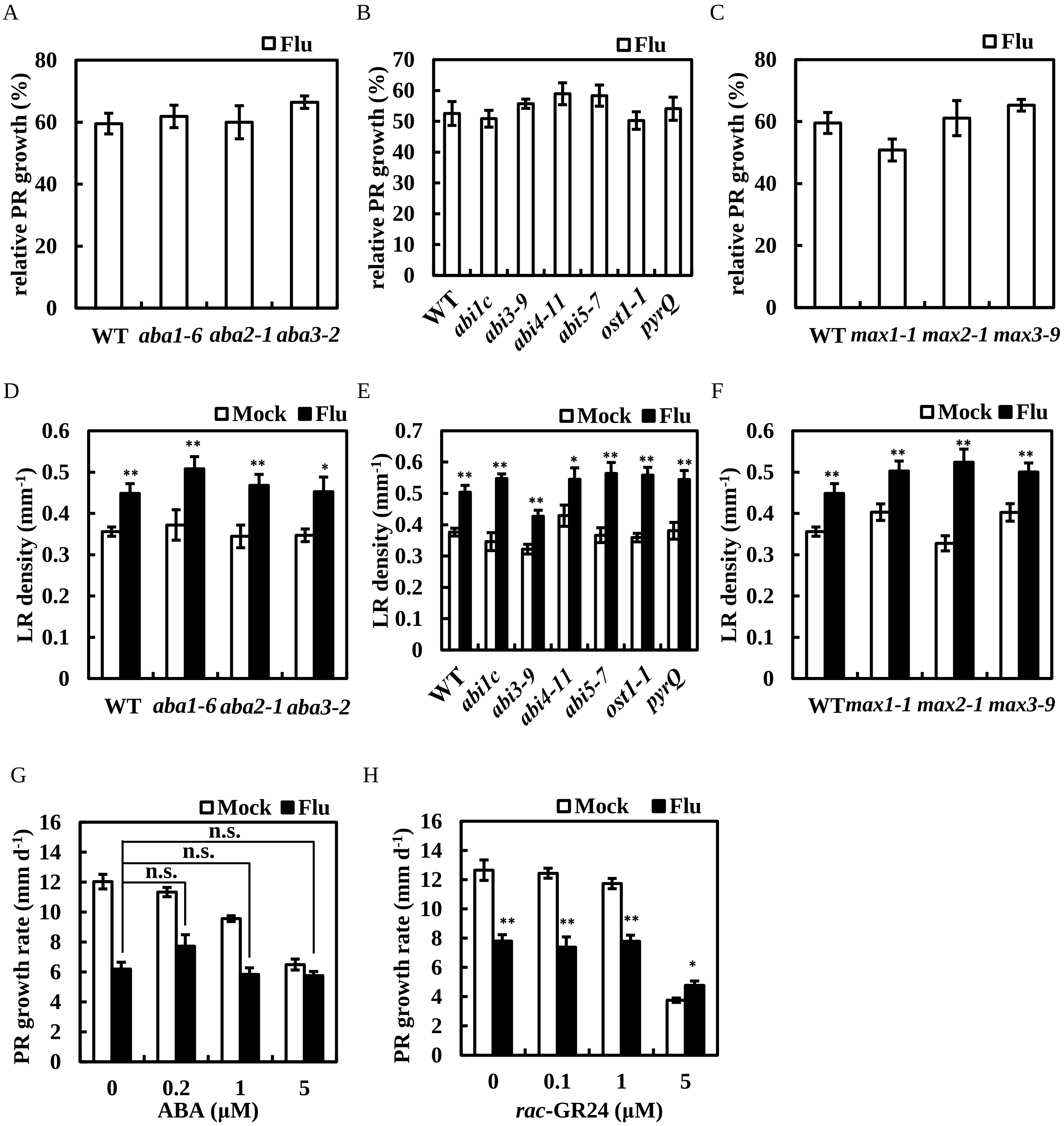


Supplemental Figure S5. ABA and SLs are not involved in fluridone-mediated regulation of root architecture. A–F) The resistance to fluridone in ABA-related and SL-related mutants. Five-day-old plants were transferred to medium with or without 800 nM fluridone. (A, D) ABA biosynthesis mutants. (B, E) ABA signaling mutants. (C, F) SL biosynthesis and signaling mutants. The relative PR growth rate compared to mock-treated plants was analyzed from 2 to 3 dat. The LR density was counted at 3 dat. G, H) Five-day-old plants were transferred to medium with or without fluridone and ABA (G) or *rac*-GR24 (H). The PR growth rate was analyzed from 2 to 3 dat. Data represent the means ± SE from 10 seedlings. *Significant differences compared to mock-treated plants in (D, E, F, H) (*: $P < 0.05$, **: $P < 0.01$; Student's *t*-test). n.s. indicates no significant difference.

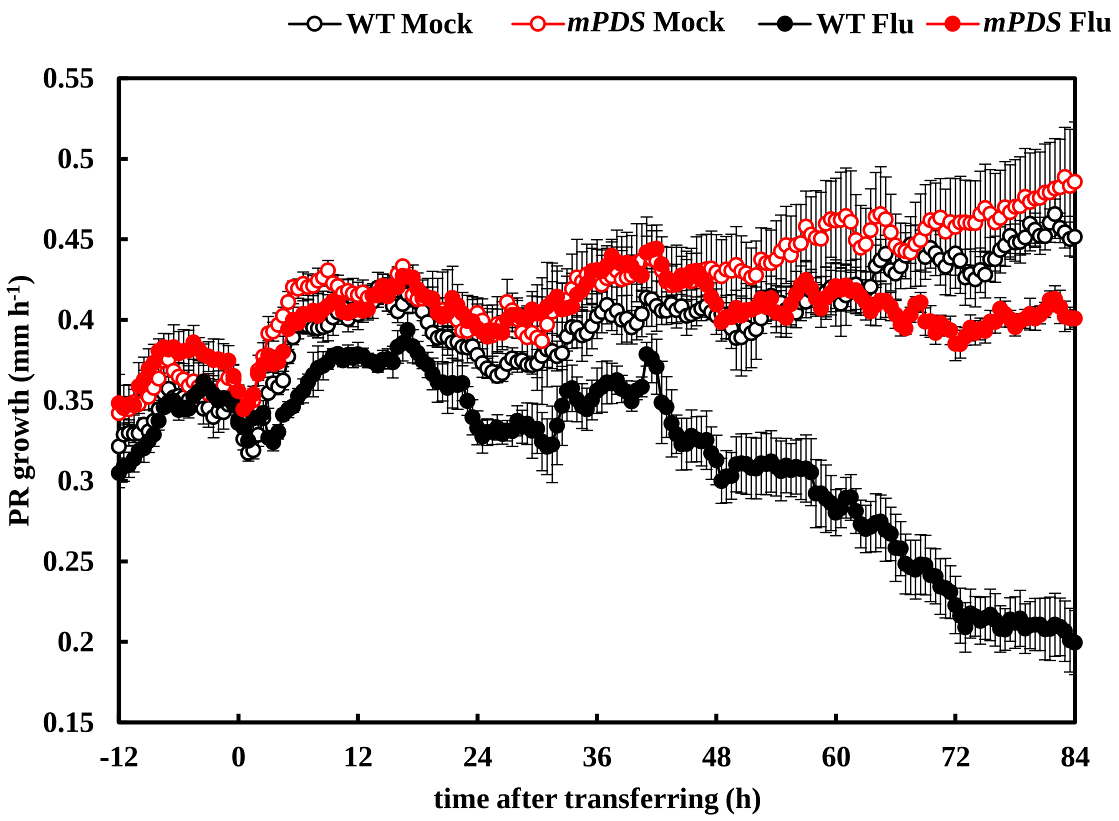


Supplemental Figure S6. The kinetic growth rate of PRs in WT and *mPDS* after fluridone treatment. Four-day-old plants of WT and *mPDS* were transferred to mock medium for a 1 d preincubation. Then plants were transferred to new medium with or without 800 nM fluridone at time 0. The PR growth rate was measured from preincubation at an interval of 30 min, which was indicated as WT mock-treated plants (hollow black), WT fluridone-treated plants (solid black), *mPDS* mock-treated plants (hollow red), and *mPDS* fluridone-treated plants (solid red). Data represent the means ± SE from 5 seedlings.

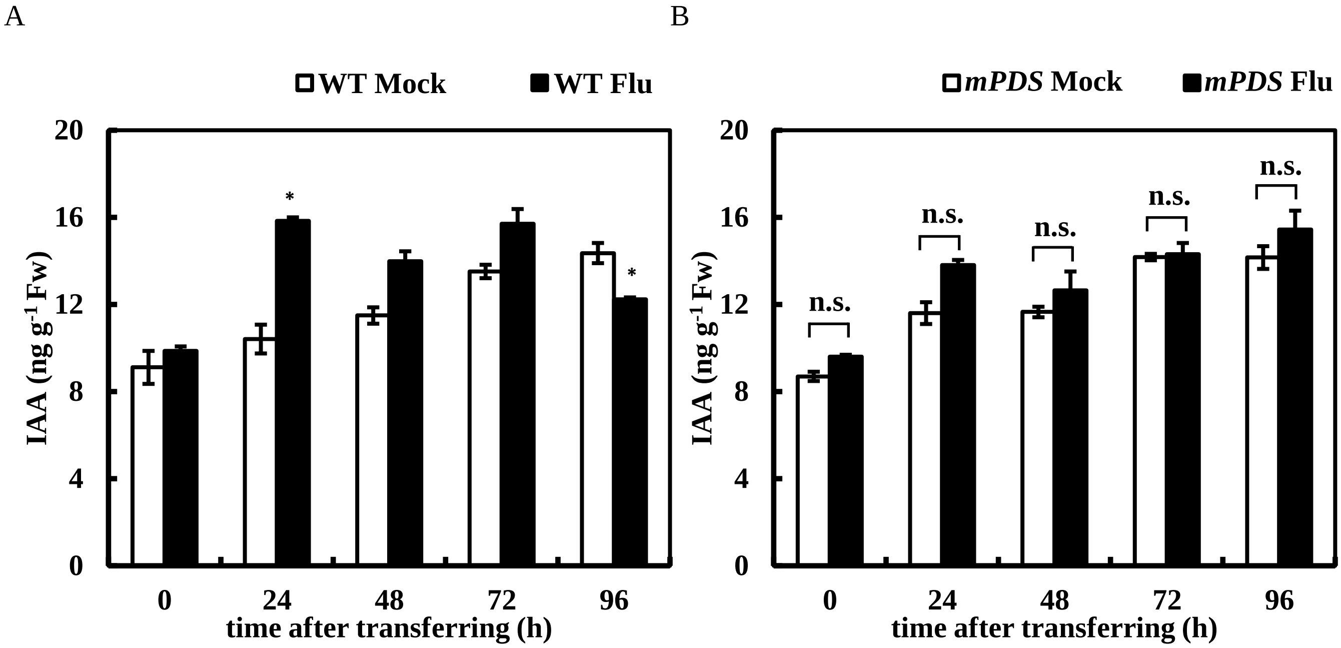


Supplemental Figure S7. Long-period change of auxin levels by fluridone. A, B) Five-day-old plants were transferred to medium with or without 800 nM fluridone at time 0. The IAA contents in the roots of WT (A) and *mPDS* (B) were measured at the indicated time points. Data represent the means ± SE from 3 biological replicates. *Significant differences compared to mock plants (*: $P < 0.05$; Student's *t*-test). n.s. indicates no significant difference.

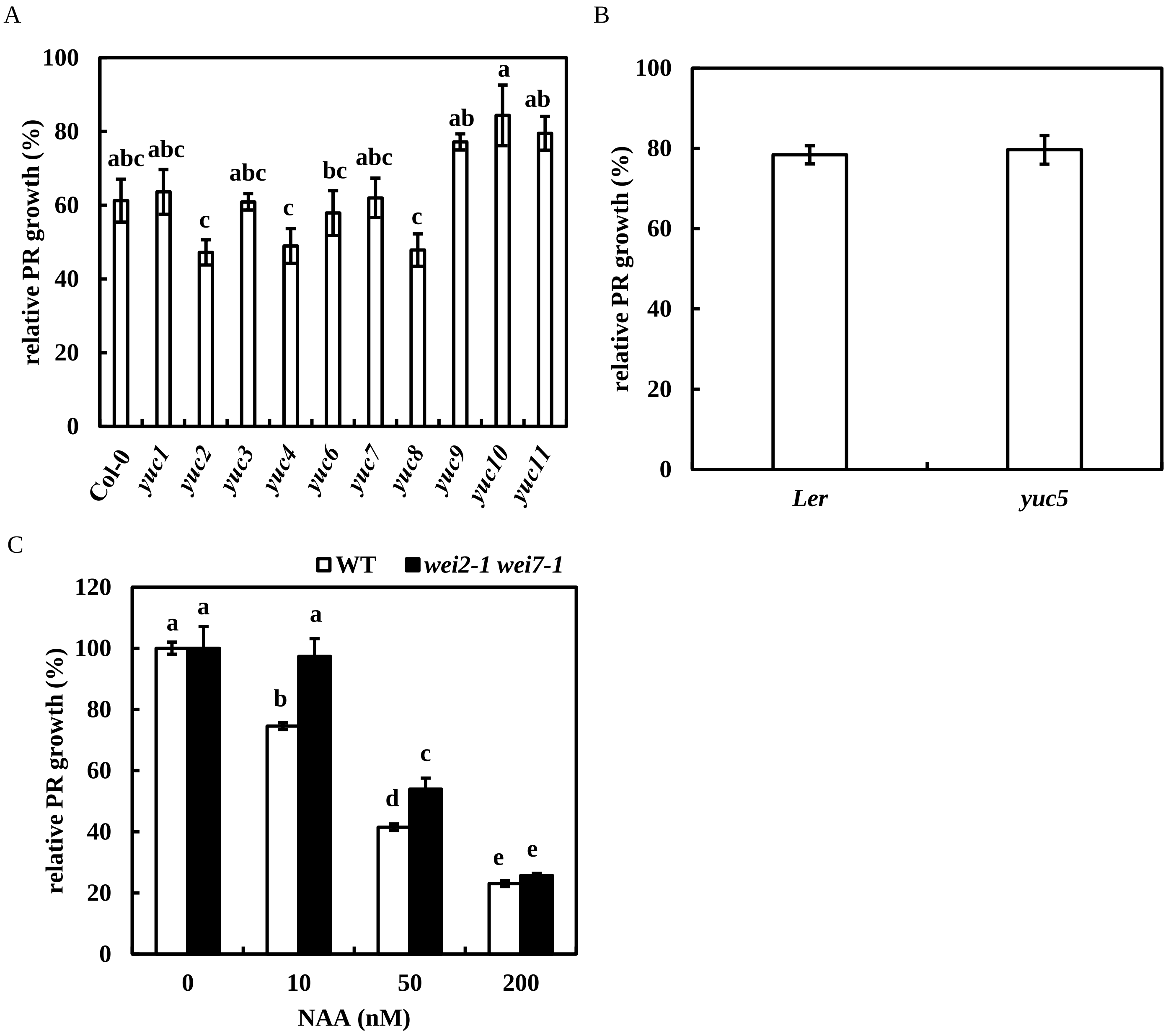


Supplemental Figure S8. The *yuc* single mutants have weak or no resistance to fluridone, and *wei2-1 wei7-1* shows resistance to low dose of exogenous NAA. A, B) Five-day-old plants were transferred to medium with or without 800 nM fluridone. (A) The *yuc* mutants in a Col-0 background. (B) The *yuc* mutant in a Landsberg *erecta* background. C) Five-day-old plants of WT and *wei2-1 wei7-1* were transferred to medium with or without NAA. The relative PR growth rate compared to mock-treated plants was analyzed from 2 to 3 dat. Data represent the means ± SE from 8-10 seedlings. Different lowercase letters above the bars in (A, C) indicate significant differences at $P < 0.05$ (one-way ANOVA following Tukey-Kramer test). n.s. indicates no significant difference.

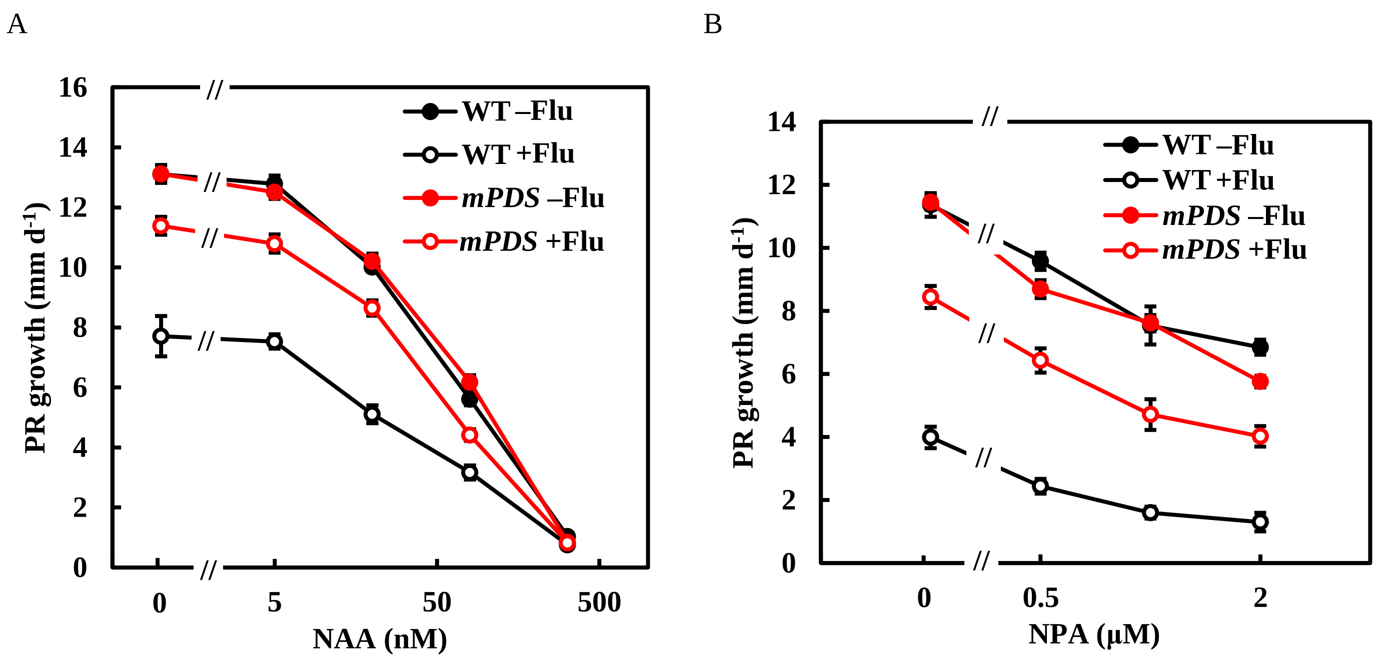


Supplemental Figure S9. Exogenous auxin and NPA suppress PR growth in an additive manner with fluridone. A, B) Five-day-old plants of WT and *mPDS* were transferred to medium with or without 800 nM fluridone and NAA (A) or NPA (B). The PR growth rate was analyzed from 2 to 3 dat in (A) or 3 to 4 dat in (B). Data represent the means ± SE from 8 seedlings. For (A), the relative PR growth of fluridone-treated plants to non-fluridone-treated plants is as follows: 58.8% (WT, 0 nM NAA), 86.8% (*mPDS*, 0 nM NAA), 58.9% (WT, 5 nM NAA), 86.3% (*mPDS*, 5 nM NAA), 51.0% (WT, 20 nM NAA), 84.7% (*mPDS*, 20 nM NAA), 47.2% (WT, 80 nM NAA), 79.9% (*mPDS*, 80 nM NAA), 72.9% (WT, 320 nM NAA) and 97.8% (*mPDS*, 320 nM NAA). For (B), the relative PR growth of fluridone-treated plants to non-fluridone-treated plants is as follows: 35.1% (WT, 0 µM NPA), 73.8% (*mPDS*, 0 µM NPA), 25.5% (WT, 0.5 µM NPA), 73.9% (*mPDS*, 0.5 µM NPA), 21.2% (WT, 1 µM NPA), 61.9% (*mPDS*, 1 µM NPA), 19.0% (WT, 2 µM NPA) and 69.8% (*mPDS*, 2 µM NPA).

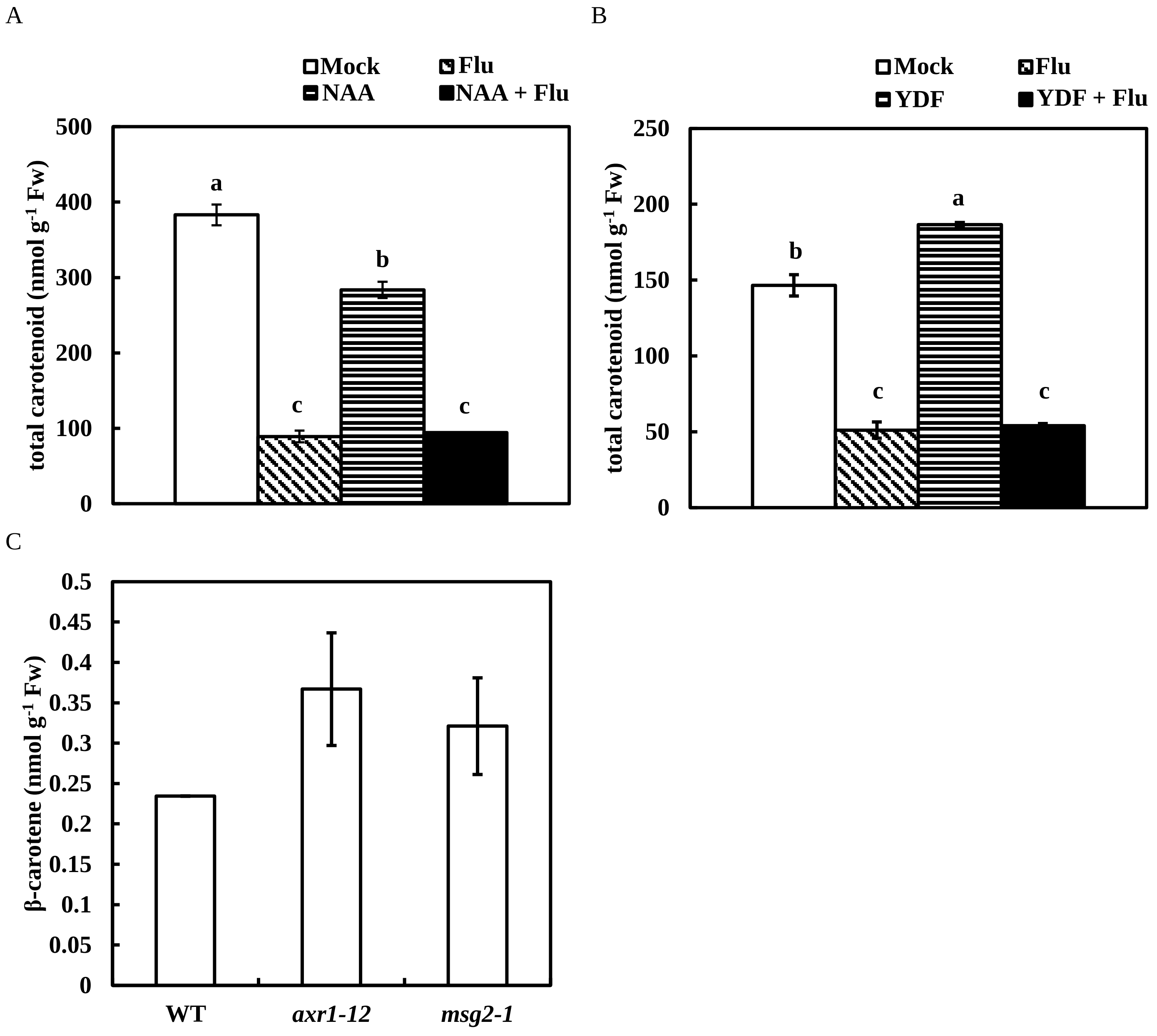


Supplemental Figure S10. Carotenoid contents in the shoots are negatively regulated by auxin, and auxin signaling mutants insignificantly increase carotenoid contents in the roots. A, B) Five-day-old plants were transferred to medium with or without 800 nM fluridone and 200 nM NAA (A) or 100 μM YDF (B). The bars indicate mock (white), 800 nM fluridone (diagonal stripe black), NAA (A) or YDF (B) (horizontal striped black) and fluridone + NAA (A) or fluridone + YDF (B) (black). The levels of total carotenoids in the shoots were measured at 4 dat in (A) and at 5 dat in (B). C) The β-carotene levels of the auxin signaling mutants in the roots were measured at 7 d after germination. Data represent the means ± SE from 3 biological replicates. Different lowercase letters above the bars in (A, B) indicate significant differences at $P < 0.05$ (one-way ANOVA following Tukey-Kramer test).

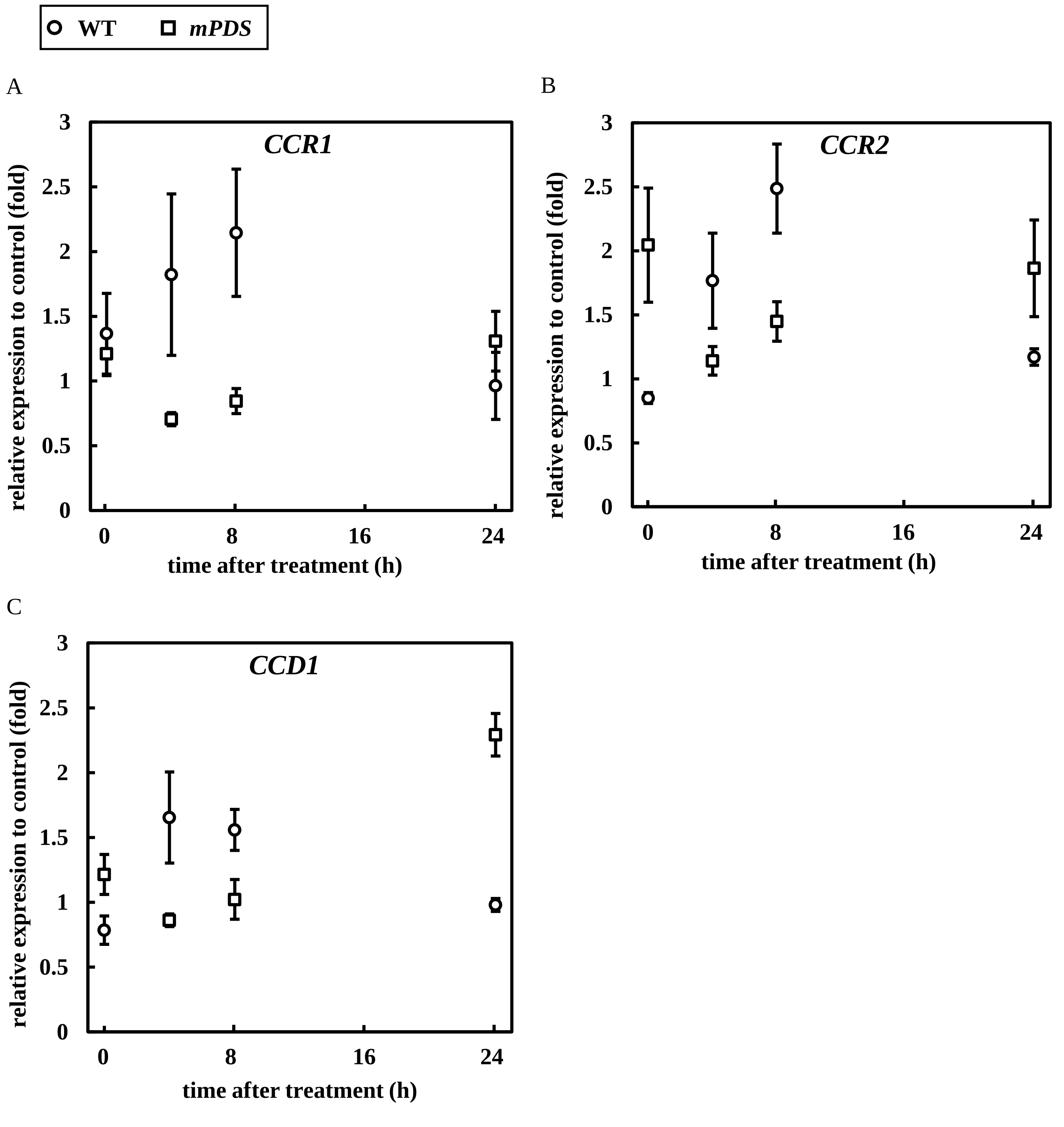


Supplemental Figure S11. The expression of *CCR1*, *CCR2* and *CCD1* is somewhat induced by fluridone. Five-day-old plants of WT and *mPDS* were transferred to medium with or without 800 nM fluridone at time 0. A–C) The relative expression of *CCR1*, *CCR2*, and *CCD1* to mock-treated plants after fluridone treatment in WT (open circle) and *mPDS* (open square). Data represent the means ± SE from 3 biological replicates.

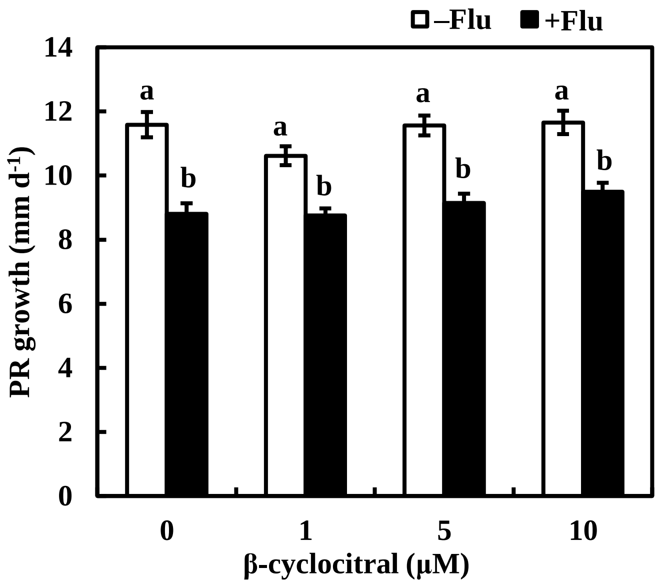


Supplemental Figure S12. β-cyclocitral is not involved in carotenoid-mediated regulation of PR growth. Five-day-old plants were transferred to medium with or without 800 nM fluridone and β-cyclocitral. The PR growth rate was analyzed from 1 to 2 dat. Data represent the means ± SE from 8 seedlings. Different lowercase letters above the bars indicate significant differences at $P < 0.05$ (one-way ANOVA following Tukey-Kramer test).

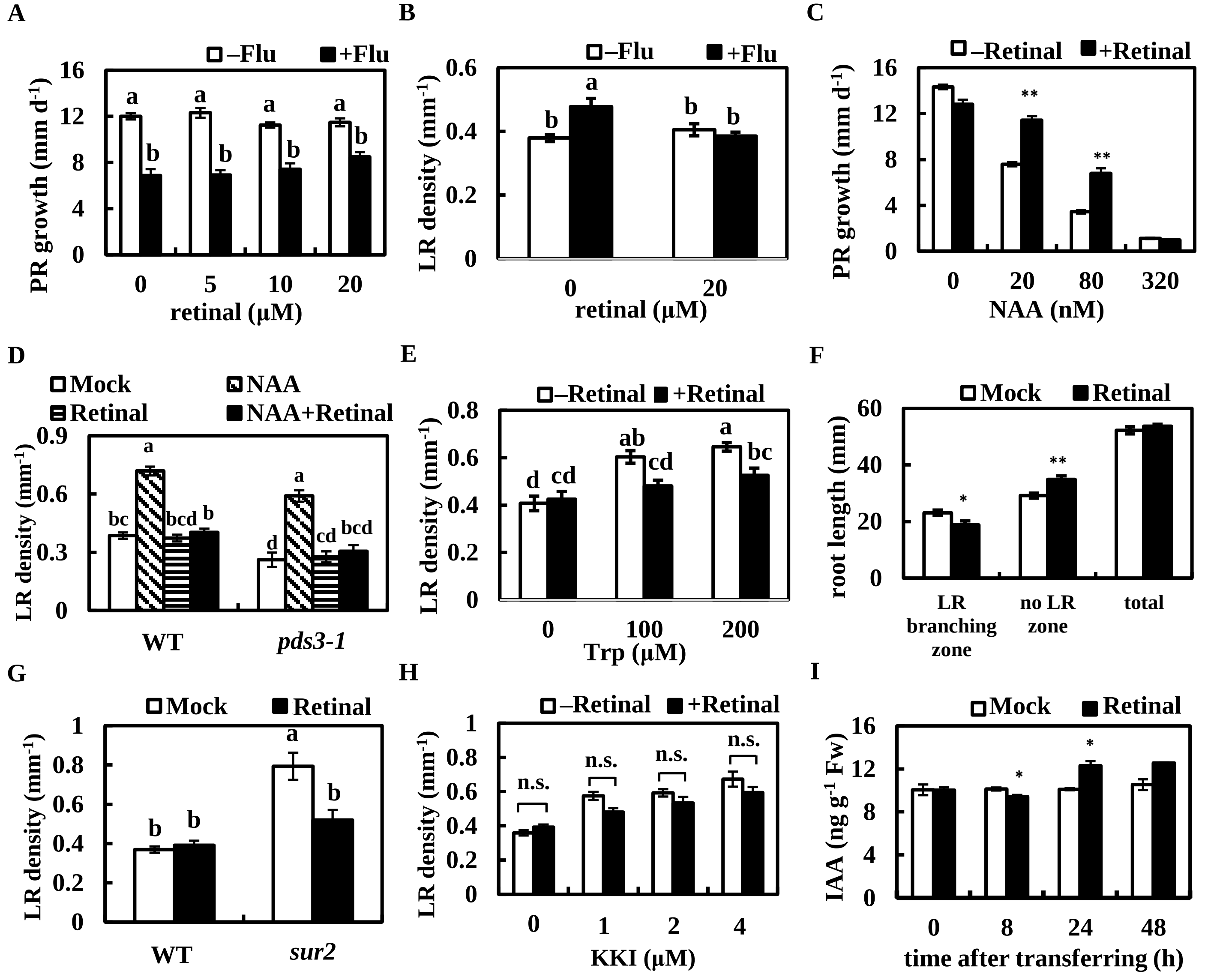


Supplemental Figure S13. Retinal is responsible for the fluridone effect and regulates auxin levels and response. A, B) Five-day-old plants were transferred to medium with or without 800 nM fluridone and retinal. C) Five-day-old plants were transferred to medium with or without 20 μM retinal and NAA. D) Five-day-old plants of WT and *pds3-1* were transferred to medium with or without 20 μM retinal and 20 nM NAA. E) Five-day-old plants were transferred to medium with or without 20 μM retinal and tryptophan. F) The length of LR branching zones (from the root–shoot junctions to the youngest LRs toward the root tips), no LR zones (from the youngest LRs to the root tips) and total roots. Five-day-old plants were transferred to medium with or without 20 μM retinal. G) Five-day-old plants of WT and *sur2* were transferred to medium with or without 20 μM retinal. H) Five-day-old plants were transferred to medium with or without 20 μM retinal and KKI. The PR growth rate in (A, C) was measured from 2 to 3 dat. The root length in (F) was measured at 3 dat. The LR density in (B, D, E, G, H) was counted at 3 dat. The bars in (D) indicate mock (white), 20 nM NAA (diagonal stripe black), 20 μM retinal (horizontal striped black) and NAA + retinal (black). I) IAA quantification by retinal treatment. Five-day-old plants were transferred to medium with or without 20 μM retinal at time 0. The IAA contents in the roots were measured at the indicated time points. Data represent the means ± SE from 8 to 10 seedlings in (A–H) and from 3 biological replicates in (I). Different lowercase letters above the bars (A, B, D, E, G) indicate significant differences at $P < 0.05$ (one-way ANOVA following Tukey-Kramer test). *Significant differences compared to non retinal-treated plants in (C, F, I) (*: $P < 0.05$, **: $P < 0.01$; Student's *t*-test). n.s. indicates no significant difference.

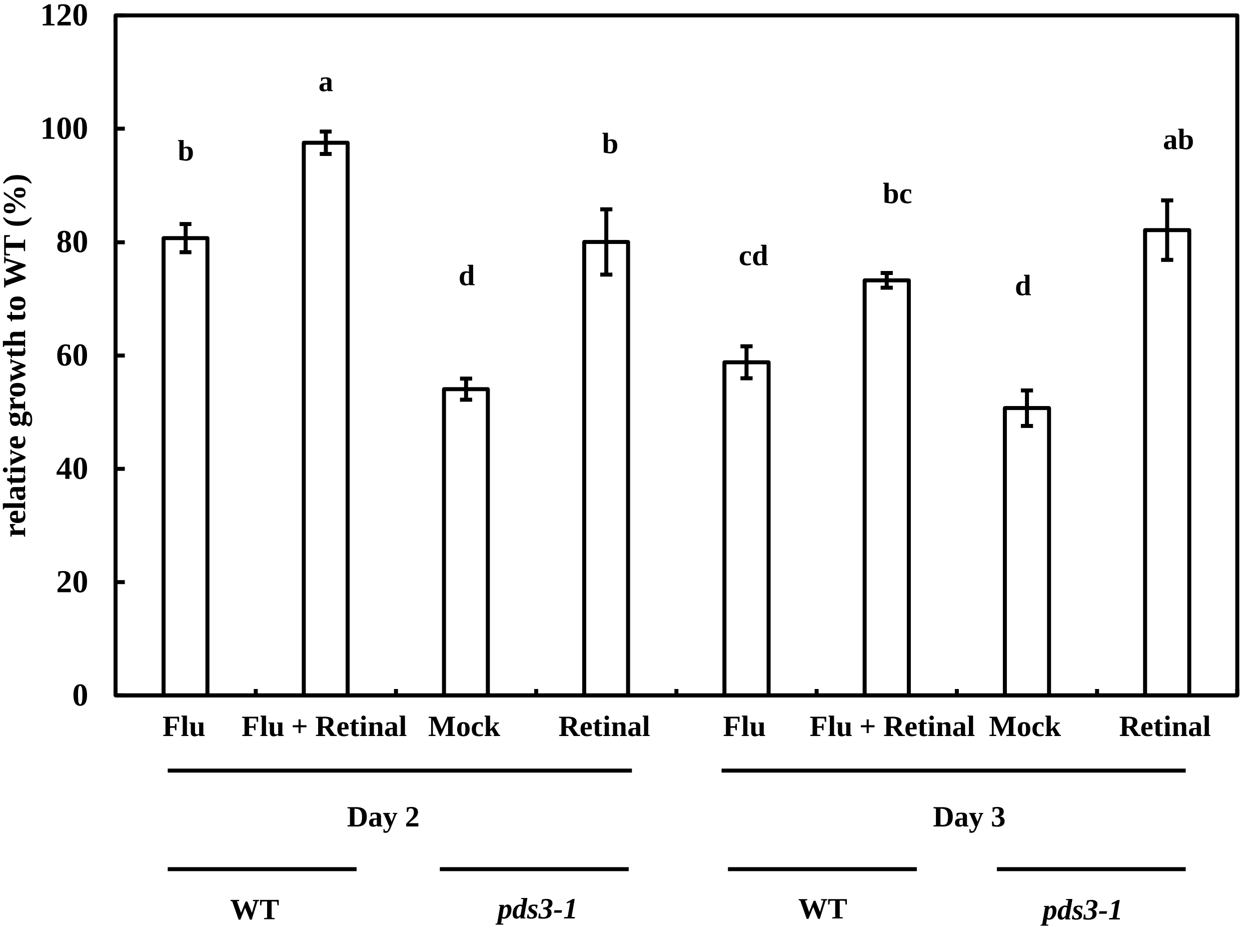


Supplemental Figure S14. The different effect of retinal to fluridone-treated WT plants and *pds3-1* plants in root growth recovery. Five-day-old plants of WT were transferred to medium in mock treatment, 800 nM fluridone treatment and co-treatment of 800 nM fluridone and 20 μM retinal. Five-day-old plants of *pds3-1* were transferred to medium with or without 20 μM retinal. The relative PR growth rate compared to the mock-treated WT plants was analyzed from 1 dat to 2 dat or from 2 dat to 3 dat. Data represents the means ± SE from three independent technical replicates. Different lowercase letters above the bars indicate significant differences at $P < 0.05$ (one-way ANOVA following Tukey-Kramer test).

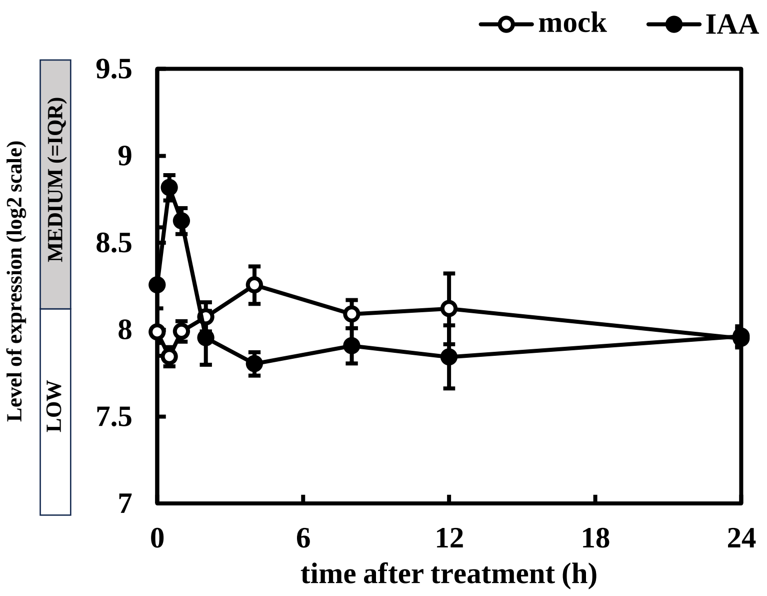


Supplemental Figure S15. *CCD7* expression is rapidly induced by exogenous IAA treatment. Public microarray data were retrieved and analyzed using GENEVESTIGATOR and the expression of *CCD7* was plotted based on Affymetrix Arabidopsis ATH1 Genome Array data (Experiment ID: AT-00655). Data are expressed as log2 scale and represent the means ± SE from 3 biological replicates.
