## Supplemental Tables 1-3 for "The exogenous application of the apocarotenoid retinaldehyde negatively regulates auxin-mediated root growth"

Supplemental Table S1. PR growth rate in mock-treated auxin-related mutants. The results were indicated by mean ± SE (n = 20 for signaling and polar transport mutants, n = 8 for the other mutants).

| Genotype | Growth rate (mm $d^{-1}$) |
|---|---|
| WT (For signaling and polar transport mutants) | 15.40 ± 0.19 |
| *msg2-1* | 16.34 ± 0.23 |
| *slr-1* | 11.74 ± 0.38 |
| *arf19-1i/nph4-1* | 12.45 ± 0.37 |
| *axr1-12* | 17.30 ± 0.33 |
| *aux1-21* | 9.30 ± 0.83 |
| WT (For *wei2*, *wei7* and *wei2 wei7* mutants) | 9.89 ± 0.42 |
| *wei2-1* | 9.99 ± 0.49 |
| *wei7-1* | 10.25 ± 0.16 |
| *wei2-1 wei7-1* | 1.60 ± 0.19 |
| WT (For *yuc* mutants) | 12.18 ± 0.34 |
| *yuc1* | 12.33 ± 0.40 |
| *yuc2* | 11.94 ± 0.25 |
| *yuc3* | 11.49 ± 0.63 |
| *yuc4* | 11.98 ± 0.63 |
| *yuc6* | 11.21 ± 0.84 |
| *yuc7* | 11.43 ± 0.50 |
| *yuc8* | 12.64 ± 0.26 |
| *yuc9* | 11.04 ± 0.55 |
| *yuc10* | 8.99 ± 0.69 |
| *yuc11* | 11.44 ± 0.50 |
| *yucQ* | 3.83 ± 0.53 |
| Ler (For *yuc5* mutant) | 10.89 ± 0.20 |
| *yuc5* | 8.93 ± 0.27 |

Supplemental Table S2. Endogenous IAA concentration in fluridone treatment in the shoots. Five-day-old plants were transferred to mock or 800 nM fluridone medium at time 0. The IAA contents in the shoots were measured at indicated timepoints. The results were indicated by mean ± SE (n = 3). The unit is ng $g^{-1}$ Fw.

| | 0 h | 8 h | 24 h |
|---|---|---|---|
| Control | 11.41 ± 1.63 | 12.92 ± 1.40 | 11.55 ± 1.25 |
| Fluridone | 11.35 ± 1.23 | 11.44 ± 1.01 | 10.86 ± 0.84 |

Supplemental Table S3. Gene-specific primers used for qRT–PCR analysis.

| Gene name | Primer name | Primer Sequence (5'-3') | Reference |
|---|---|---|---|
| *ACTIN2* | ACTIN2-27S | 5'-CGCTCTTTCTTTCCAAGCTCATA-3' | Xu et al., 2017 |
| | ACTIN2+55 AS | 5'-CCATACCGGTACCATTGTCACA-3' | |
| *IAA19* | IAA19+390S | 5'-CTTCGGTTTCCGTGGCATCG-3' | Xu et al., 2017 |
| | IAA19+521 AS | 5'-CATGACTCTAGAAACATCCC-3' | |
| *YUC9* | YUC9+542S | 5'-ATAAGTCCGGCGAGAAATTCAGAG-3' | Xu et al., 2017 |
| | YUC9+682 AS | 5'-TCGGTAAAACATGAACCGAG-3' | |
| *CCR1* | CCR1+3599 S | 5'-AAGAGCAAGCGTTTGACAGGTAAG-3' | This study |
| | CCR1+3678 AS | 5'-TACGATTGCGATGAAGGAACTGG-3' | |
| *CCR2* | CCR2+2888 S | 5'-CATCGGGTAGCTGCTGATATTGGG-3' | This study |
| | CCR2+2968 AS | 5'-AGCCAACCAAGTAAACCAAGAAGG-3' | |
| *CCD1* | CCD1+2574 S | 5'-TATGTTCCGCGTGAGACAGCAG-3' | This study |
| | CCD1+2665 AS | 5'-TCACAGTCACGCATGATTTCCC-3 | |
| *CCD7* | CCD7+2077 S | 5'-CGTTGGTGAGCCCATGTTTGTC-3' | This study |
| | CCD7+2180 AS | 5'-TCTCTCCACCGAAACCGCATACTC-3' | |
| *CCD8* | CCD8+2715 S | 5'-GTGCAACCCATGAGGATGATGGAG-3' | This study |
| | CCD8+2839 AS | 5'-CCATAGGGAAACTTGGCTCTTGC-3' | |

**Xu D, Miao J, Yumoto E, Yokota T, Asahina M, Watahiki M** (2017) *YUCCA9*-mediated auxin biosynthesis and polar auxin transport synergistically regulate regeneration of root systems following root cutting. *Plant and Cell Physiology* **58:** 1710-1723
