## Supplemental Movie 1 Legend for "The exogenous application of the apocarotenoid retinaldehyde negatively regulates auxin-mediated root growth"

### Supplemental Movie Legend

Movie S1 The *DR5* bioluminescence video during fluridone treatment of the WT and *mPDS* plants. Five-day-old plants were transferred to new medium with or without 800 nM fluridone. Scale bar = 5 mm.
